# Mucinolysome in gut microbiomes of farm animals and humans

**DOI:** 10.1101/2025.10.14.682383

**Authors:** Jerry Elorm Akresi, Thi Van Thanh Do, Zhisong Cui, N.R. Siva Shanmugam, Sarah Moraïs, Itzhak Mizrahi, Edward A Bayer, Jennifer Auchtung, Yanbin Yin

## Abstract

Mucins are glycoproteins that create a protective barrier protecting host tissues from microbial pathogens and are instrumental for host health. Here, we provide evidence that mucin glycan degradation in the gut can be mediated by mucinolysomes, defined as extracellular multi-enzyme complexes specializing in mucin glycan degradation. We computationally predicted the presence of mucinolysomes across 63 metagenome-assembled genomes (MAGs) and two isolated genomes of anaerobic *Limousia* bacteria, including seven MAGs from human samples of six countries. All 65 genomes were found to display core mucinolysome components, consisting of 3∼6 scaffoldins (containing up to 12 cohesin modules) and up to 22 dockerin-containing mucin glycan-degrading CAZymes (carbohydrate active enzymes). The organization of mucinolysomes allows the assembly of up to 24 CAZymes in the same complex. We validated that a cultivated *Limousia* strain ET540 from chicken cecum can support growth on mucins as its sole carbon source, triggering the expression of most mucinolysome-related genes, including both scaffoldins and CAZymes. We also modeled the assembly of proteins into a multi-enzyme complex by predicting the cohesin-dockerin interactions among most of the mucinolysome proteins using AlphaFold3. While mucinolysosome-encoding *Limousia* have low abundance in different animal hosts, their abundance and prevalence are higher in farm animals than in humans, highlighting a potentially important role in livestock gut ecosystems. Our findings reveal a novel mechanism of mucin glycan degradation and provide a framework to explore microbial contributions to gut health and host-microbe interactions across species.

## INTRODUCTION

Mucins are glycoproteins that form the mucus layers on epithelial cell surfaces, providing an important physical barrier that functions to separate the epithelium from potential threats, including those of gut microbes (*1*). Excessive mucin glycan degradation by commensal and pathogenic bacteria can contribute to different gastrointestinal tract (GI) diseases (*2–4*) in humans and animals. As a result, mucin glycan degradation must be tightly regulated, as the recycling of mucin-derived glycans is essential for the nutrient cycle and microbial cross-feeding processes that sustain a stable and healthy gut ecosystem.

The human colonic mucus is mainly composed of the large mucin glycoprotein MUC2. MUC2 contains 20% proteins and 80% glycans with over 3,300 terminal sugar residues (*3*). These glycans are constructed from five different monosaccharides, L-fucose, N-acetyl-D-galactosamine, N-acetyl-D-glucosamine, D-galactose, and sialic acid (N-acetyl-neuraminic acid), that can be connected by over a dozen different glycosidic linkages in mucins (*3*). Their degradation requires a broad repertoire of carbohydrate-active enzymes (CAZymes), spanning numerous glycoside hydrolase families, including GH2, GH20, GH29, GH33, GH84, and GH95, among others, encoded in many different gut microbes (*5*). Mucins also contain non-sugar modifications, for example, with sulfate and acetyl groups, that can be removed by microbial activity. Reflecting this chemical complexity, a recent report described the extensive CAZyme arsenal from the mucinolytic bacterium, *Akkermansia muciniphila,* capable of complete degradion of O-linked mucin glycans (*6*).

Mucin-glycan degrading CAZymes are often clustered into polysaccharide utilization loci (PULs) for synergistic mucin degradation in genomes of Gram-negative gut bacteria, such as *Bacteroides thetaiotaomicron* (*7–10*). Here, we report the first description of a mucinolysome, a novel cellulosome-like system comprising multiple CAZymes, uniquely encoded by a group of Gram-positive gut bacteria, to degrade mucin glycans in the GI tracts of farm animals and humans. Cellulosomes are multienzymatic complexes, first discovered in 1983 in *Clostridium thermocellum* from hot springs, which are capable of efficient degradation of plant polymers, such as cellulose, hemicellulose and other plant cell wall polysaccharides (*11, 12*). Cellulosomes contain hallmark protein domains, cohesins (Coh) and dockerins (Doc), which are key structural modules that interact to form Lego-like multi-protein complexes (*13*). Cohesins are typically found as tandem repeats in scaffoldins, large structural proteins that generally lack catalytic functions. Dockerins, on the other hand, are present in CAZymes such as cellulases and hemicellulases. The cohesin-dockerin (Coh-Doc) interaction defines incorporation of the enzymes into the complex and cellulosome formation. Scaffoldins provide sites for docking of multiple CAZymes to form the highly organized swissknife-like multi-enzyme cellulosome complexes for the coordinated degradation of complex carbohydrates (*14*). More recently, the “amylosome”, a cellulosome-like system, was discovered in *Ruminococcus bromii* from human gut for resistant starch degradation (*15, 16*), suggesting that the cellulosome paradigm may also be extended to other complex carbohydrate degradation, such as mucin glycans.

In this study, we extensively searched existing human and animal gut metagenomic databases and found a total of 65 gut bacterial genomes that may encode mucinolysomes. These genomes include 63 metagenome assembled genomes (MAGs) and two bacterial isolate genomes, which were isolated from chicken caecum (*17*). One of the isolated genomes (GCF_002160515.1, *Anaeromassilibacillus* sp. An172) was previously published in draft status (*17*), and the other (strain ET540) was sequenced and assembled into a gapless complete genome in this study. The 63 MAGs included seven from humans of six different countries and 56 from farm animals. Using the genome taxonomy database (GTDB) (*18*) we assigned all 65 genomes to a newly defined genus *Limousia* (*19*). Using a single copy core gene phylogeny, we classified these genomes into three distinct species. We experimentally verified that the *Limousia* strain ET540 grew well with mucin as a sole carbon source, and generated RNA-seq data comparing gene expression in defined medium with glucose (control) and with mucin. We observed an upregulation of a majority of mucinolysome genes in ET540 grown on mucins. We also computationally predicted Coh-Doc interactions in the mucinolysome system using AlphaFold3. Finally, following our previous metagenomic read mapping approach (*20*), we investigated the relative abundance and prevalence of these mucinolysome-encoding *Limousia* species in humans and farm animals. We found that mucinolysome-encoding *Limousia* were detected in low abundance across different animal hosts, with prevalence and abundance higher in farm animals than in human samples.

## RESULTS

### Seven human MAGs are identified to encode mucinolysomes

The co-presence of dockerin and cohesin modules in the same genome signifies the existence of cellulosome-like protein complexes in a bacterium. From 289,232 genomes of the Unified Human Gastrointestinal Genome (UHGG) database (*21*) (**Figure 1A**), we found 4,333 (1.5%) MAGs that encode both dockerins and cohesins (**Figure 1B**). As expected, most of these MAGs are from the Firmicutes phylum, and the remaining 78 (1.8% of 4,333) are from Bacteroidetes. Interestingly, human populations with a higher intake of fiber-rich diets, according to the Global Dietary Database (*22*), have higher percentages of MAGs with Coh+Doc modules (**Figure 1C**, e.g., Africa, South America), suggesting that fiber-rich diets select cellulosomal bacteria.

**Figure 1:**
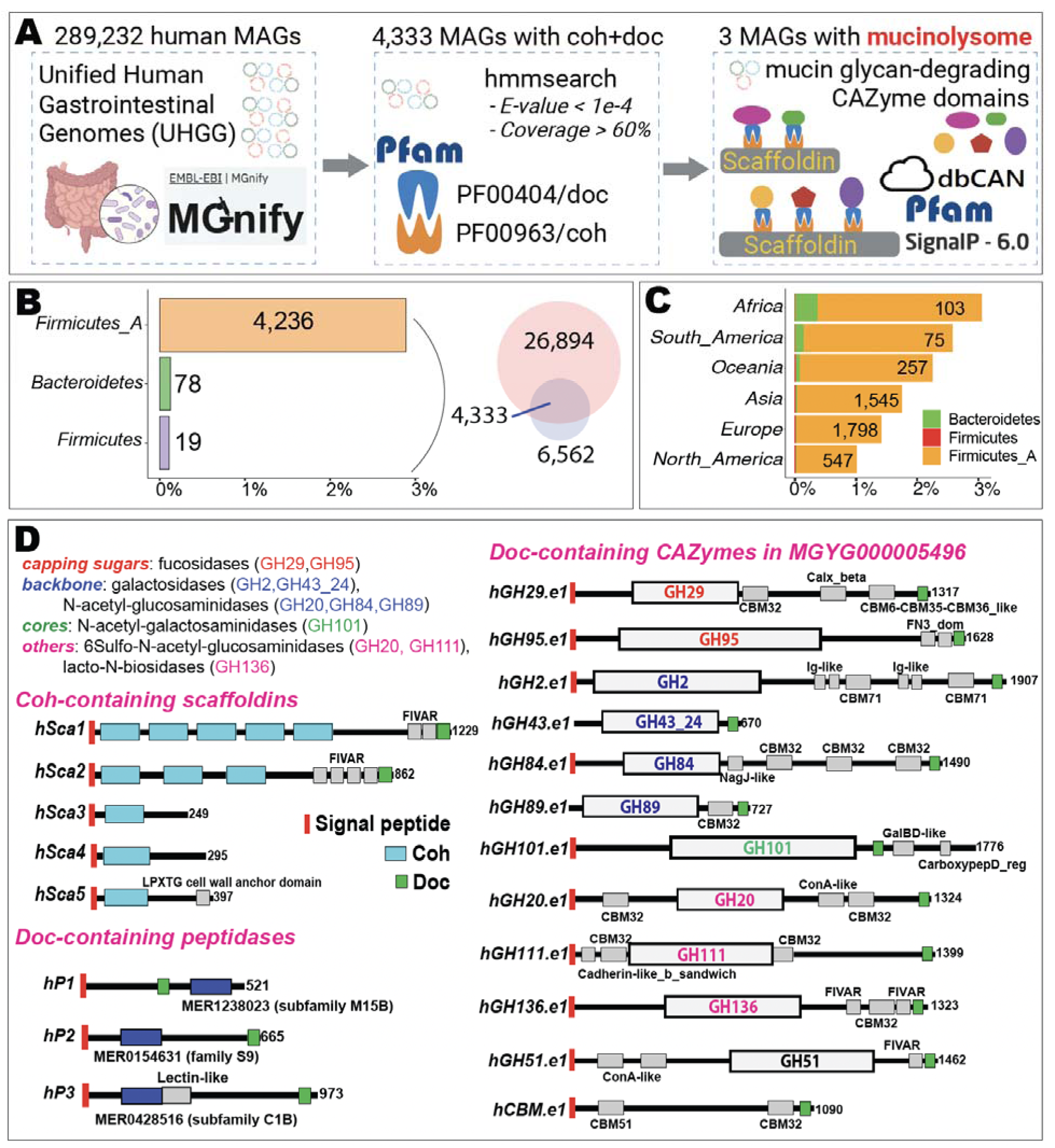
Discovery of mucinolysome-encoding MAGs in human gut microbiome. (**A**) Workflow to identify potential mucinolysomes in human gut MAGs. (**B**) The Venn diagram shows the numbers of genomes with Doc-containing proteins (26,894) and with Coh-containing proteins (6,562). The barplot shows the bacterial phyla of the 4,333 MAGs with Coh+Doc modules. The x-axis indicates the percentage of MAGs (# of MAGs with Coh+Doc / total # of MAGs in the specific phylum of UHGG). (**C**) The stacked bar plot shows the continents of the 4,333 MAGs with Coh+Doc modules. The x-axis indicates the percentage of MAGs (# of MAGs with Coh+Doc / total # of MAGs in the specific continent of UHGG). (**D**) The dbCAN/Pfam domain diagrams of Doc-containing and Coh-containing proteins in one specific UHGG MAG: MGYG000005496 (Table 1). For Doc-containing proteins, only those with CAZyme and peptidase domains are shown. CAZyme domains are colored based on their biochemical activities from the literature (2, 23, 24) and the CAZy website (cazy.org) (25).

At least 18 CAZyme families have been experimentally characterized in different gut bacteria for mucin glycan degradation (*2, 23, 24*) (**Table 1**, **Figure 1A**). In the present study, three out of the 4,333 UHGG MAGs were considered to possess mucinolysomes as they encoded both (i) scaffoldins bearing multiple Coh domains and (ii) mucin-glycan degrading CAZymes with Doc domains. One of the three MAGs (MGYG000005496, **Figure 1D**) has five scaffoldins (two with multiple Coh domains), 12 Doc-containing CAZymes, and three Doc-containing peptidases. Eleven of the 12 Doc-containing CAZymes in this MAG contain GH (glycoside hydrolase) domains. All but one of these mucinolysome proteins contain signal peptides, suggesting that they are secreted, consistent with cellulosomal systems.

**Table 1:** Seven human MAGs with mucinolysome systems.

| MAG ID (source DB) | SRA ID | Country | # of contigs | Genome length (kb) | Total # of proteins | # of Coh proteins | # of Doc proteins (*) | Disease |
| --- | --- | --- | --- | --- | --- | --- | --- | --- |
| MGYG000005496 (EBI MGnify) | ERR321631 | Denmark | 93 | 2,022,232 | 1,827 | 5 | 10 | obesity |
| MGYG000126411 (EBI MGnify) | ERR209841;<br>ERR209842;<br>ERR210693;<br>ERR210694 | Spain | 394 | 1,611,060 | 1,463 | 5 | 8 | healthy |
| MGYG000000924 (EBI MGnify) | ERR1620320 | China | 57 | 2,081,481 | 1,863 | 4 | 15 | crohn's disease |
| GCA_036306305.1 (NCBI GenBank) | SRR24600132;<br>SRR24599998;<br>SRR24600064;<br>SRR24600036 | USA | 73 | 2,139,278 | 1,896 | 4 | 20 | NA |
| GCA_019417085.1 (NCBI GenBank) | DRR171913 | Japan | 367 | 1,914,663 | 1,811 | 2 | 15 | NA |
| GCA_938048255.1 (NCBI GenBank) | ERR321631 | Denmark | 96 | 2,057,936 | 1,858 | 5 | 16 | obesity |
| GCA_958448175.1 (NCBI GenBank) | ERR9578163 | Germany | 114 | 1,969,338 | 1,892 | 6 | 11 | Alzheimer |

\* Doc-containing proteins predicted to have mucin-targeting GH domains<sup>13</sup>

Ten GH families (**Figure 1D**) in the MGYG000005496 mucinolysome have known biochemical activities in cleaving the capping sugars: fucosidases (GH29, GH95), breaking down glycan backbones: galactosidases (GH2, GH43_24), N-acetyl-glucosaminidases (GH20, GH84, GH89), degrading glycan cores: N-acetyl-galactosaminidases (GH101), and breakding down other major components in mucins: 6-Sulfo-N-acetyl-glucosaminidases (GH20, GH111) and lacto-N-biosidases (GH136). The GH43_24 family has been recently characterized as a β-galactosidase in *Akkermansia muciniphila* for mucin degradation (*6*). GH51 and GH111 have not been previously shown to degrade mucins. However, GH51 was found in a PUL in human gut *B. thetaiotaomicron* (BT3092-BT3103). This PUL contains GH2, GH43, and multiple sulfatases (*26*), suggesting GH51 in this PUL may also function in mucin glycan degradation. GH111, a keratan sulfate hydrolase family, contains an experimentally characterized 6-Sulfo-N-acetyl-glucosaminidase (*27*). Keratan sulfate has not been found in gut epithelial tissues, but is structurally similar to mucins (*28*).

To expand our search for mucinolysome MAGs, we further searched the five scaffoldin sequences present in the three MAGs against all human gut bacterial genomes in the RefSeq database. Using this approach, we found four additional human MAGs (**Table 1**) in GenBank that may also encode mucinolysomes. One of them (GCA_938048255) appears to be derived from the same metagenome sample (ERR321631) as the UHGG MGYG000005496; however, they differ in the number of contigs, genome length, and number of proteins (**Table 1**). Overall, the original microbiome samples of the seven human MAGs were sourced from both diseased and healthy human individuals of six different countries in Europe, Asia, and North America. This suggests that these mucinolysome-encoding gut bacteria are widely spread in different human populations and unlikely to be contaminations from other sources. Although, the living environments or lifestyles of these humans are unknown, one MAG (GCA_036306305.1) was published in a recent study, where microbiomes of farmers and non-farmers were compared (*29*); this specific MAG was indicated from a non-farmer (NCBI biosample ID: SAMN38021763), reinforcing our assumption that these MAG are not contaminants from animals.

### Two bacterial isolate genomes and 56 MAGs of the genus *Limousia* from farm animals also encode mucinolysomes

As one of the five scaffoldins, hSca1 (Sca1 of human MAG MGYG000005496) is 1,229 aa long with five Coh repeats (**Figure 1D**), we selected its sequence to search against the NCBI nr database and found that hSca1 has a very similar (98.94% sequence identity) protein hit (WP_087376890.1) in *Anaeromassilibacillus sp.* An172. The An172 genome (GCF_002160515.1) was sequenced from a pure bacterial culture isolated from chicken caecum (*17*). Here, we sequenced ET540, a closely related strain to An172, which was also isolated from chicken caecum. We generated the complete genome of ET540 with no gaps, annotated the two isolate genomes (ET540 and An172), and confirmed that they contained all the mucinolysome-related genes that the human MGYG000005496 possesses (**Figure S1**).

We further expanded the search for mucinolysomes to additional MAG databases. These databases included MAGs from humans living in different lifestyles (*29, 30*), from different farm animals (cows (*31, 32*), pigs (*32–34*), chickens (*35, 36*), sheep (*37*)), and from the Global Microbial Gene Catalog (GMGC) (*34*). Using the same workflow (**Figure 1A**), we found 56 additional mucinolysome-encoding MAGs (**Table S1**) from different animal hosts, including 40 from chickens, seven from pigs, and nine from cows (**Figure 2A**, column 3). These MAGs were built from metagenomes sampled from Europe (France, Germany, Spain, Denmark, Czech Republic), North America (USA, Canada), Asia (China, Japan), and Africa (Ethiopia) (**Figure 2A**, column 2). All these MAGs contain 3∼6 scaffoldins (with up to 12 Coh domains, **Figure 2A**, columns 4 and 7) and up to 22 Doc-containing CAZymes with mucin-degrading GH domains (**Figure 2A**, column 5). In addition, dockerins were also found in up to 31 other proteins with peptidase or other predicted functional domains (**Table S2, Figure 2A**, column 6). The An172 genome (GCF_002160515.1) encodes the most (52) Doc domains.

**Figure 2:**
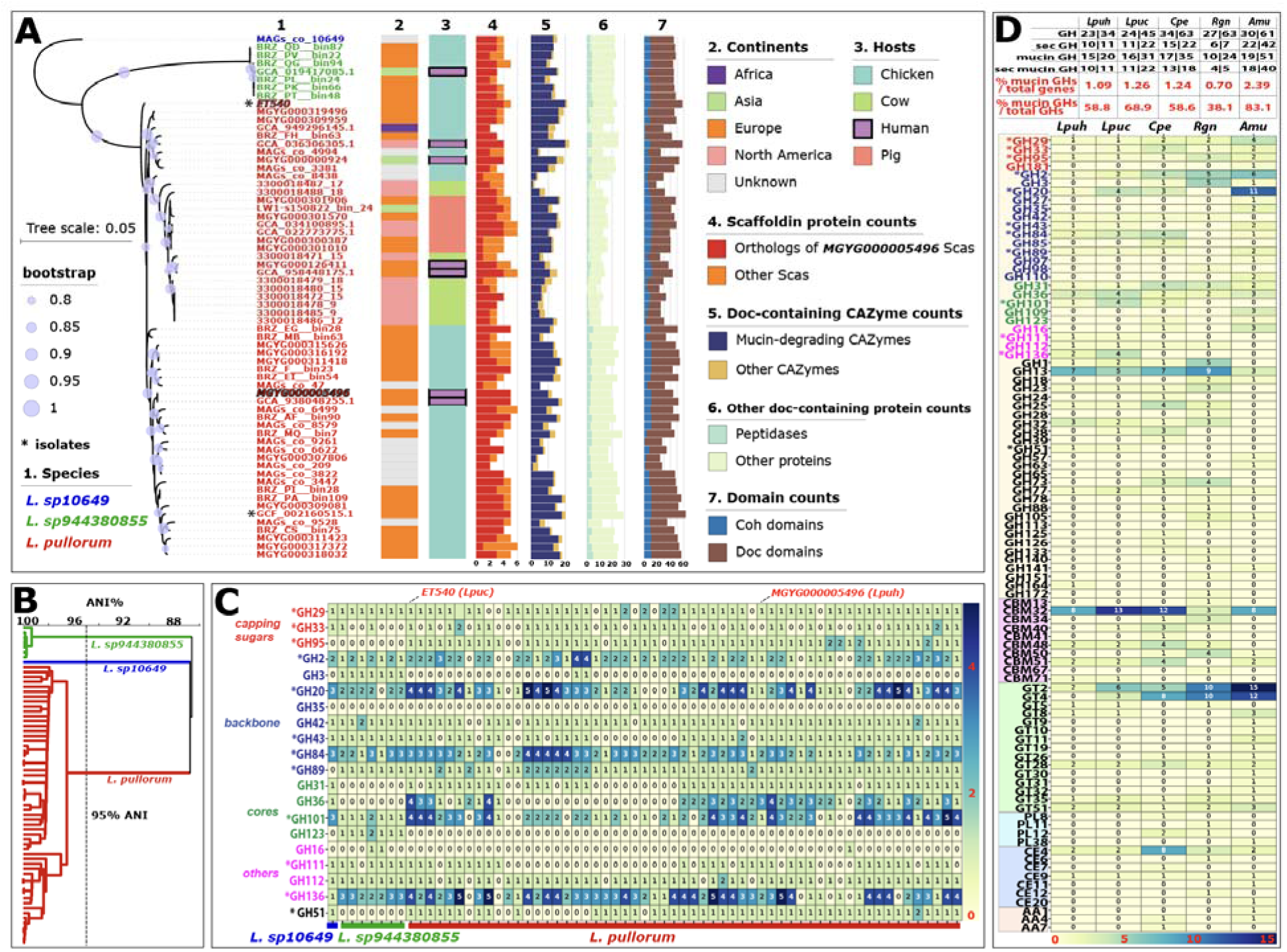
Sixty-five Limousia genomes and mucin-glycan degrading CAZymes. (**A**) Phylogeny of the 65 genomes (see **Table S1** and **Table S2** for more info) built from the single copy core gene alignment. Two genomes are isolate genomes (GCF_002160515.1 and ET540, indicated with *), and the rest are MAGs. Only ET540 has a complete genome without gaps. All others are in contig or scaffold status. MGYG000005496 and ET540 are highlighted with italicized and stroked fonts. Three species are identified according to this phylogeny. (**B**) Average nucleotide identity (ANI) dendrogram of the 65 genomes. (**C**) Heatmap of mucin glycan-degrading GH families (rows) in the 65 genomes (columns). Numbers are the protein counts of each family, irrespective of whether the proteins are Doc-containing or not. Families present in ET540 mucinolysome (Figure 3) are indicated with *. (**D**) Heatmap of all CAZyme families (rows) in two Limousia genomes and three known mucin glycan-degrading genomes (columns). Numbers are the protein counts of each family. Colored GH families (26 in total) are known for mucin glycan degradation according to the literature. The table on the top shows the family count and protein count separated by “|”. Counts for all GHs, secreted GHs, all mucin glycan GHs, and secreted mucin glycan GHs are listed (details in **Table S3**). The bottom two columns are the % of mucin glycan-degrading GHs calculated based on different denominators. The first one is calculated as the count of mucin glycan-degrading GHs divided by the total gene count in the genome. The second one is calculated as the count of mucin glycan-degrading GHs divided by the total GH count in the genome. Lpuh: Limousia pullorum MGYG000005496, Lpuc: Limousia pullorum ET540, Cpe: Clostridium perfringens ATCC 13124, Rgn: Ruminococcus gnavus ATCC 29149, Amu: Akkermansia muciniphila ATCC BAA 835.

Altogether, 65 genomes (63 MAGs and two isolate genomes, **Figure 2A**) were found to potentially encode mucinolysomes (**Figure S1**). All genomes were taxonomically annotated as *Anaeromassilibacillus* or *Acutalibacteraceae* sp. in their original databases. By running these genomes through GTDB v220 (*18*), we classified all of them to a newly defined genus, *Limousia*, and most were assigned to the species *Candidatus Limousia pullorum* (*38*). A phylogenic tree was built from the single copy core protein sequence alignment of these 65 genomes (**Figure 2A**). We identified three distinct species, which were also supported by a genome average nucleotide identity (ANI) analysis using 95% as the threshold (**Figure 2B**). The phylogeny and ANI dendrogram also support that the *L. pullorum* genomes can be further divided into two subspecies, each of which contains an isolate genome (An172 and ET540). Following this new GTDB classification, we now renamed all these formerly known *Anaeromassilibacillus* genomes as *Limousia* genomes.

### *L. pullorum* ET540 from chicken has a more complex mucinolysome system than MGYG000005496 from human and *Clostridium perfringens*

Compared to the human MAG MGYG000005496 (**Figure 1D**), the chicken ET540 genome has a more complex mucinolysome system (**Figure 3A**). ET540 encodes 21 Doc-containing CAZymes of 12 GH families, including a GH33 sialidase missing in MGYG000005496. ET540 has four GH101, four GH20, three GH136, and two GH84 proteins, compared to one copy of each in MGYG000005496. Interestingly, one of the GH20 proteins also contains a peptidase domain. Therefore, ET540 has a total of six Doc-containing peptidases (**Figure 3A**), compared to three in MGYG000005496. All scaffoldins and all Doc-containing peptidases comprise a signal peptide, in a manner similar to most Doc-containing CAZymes.

**Figure 3:**
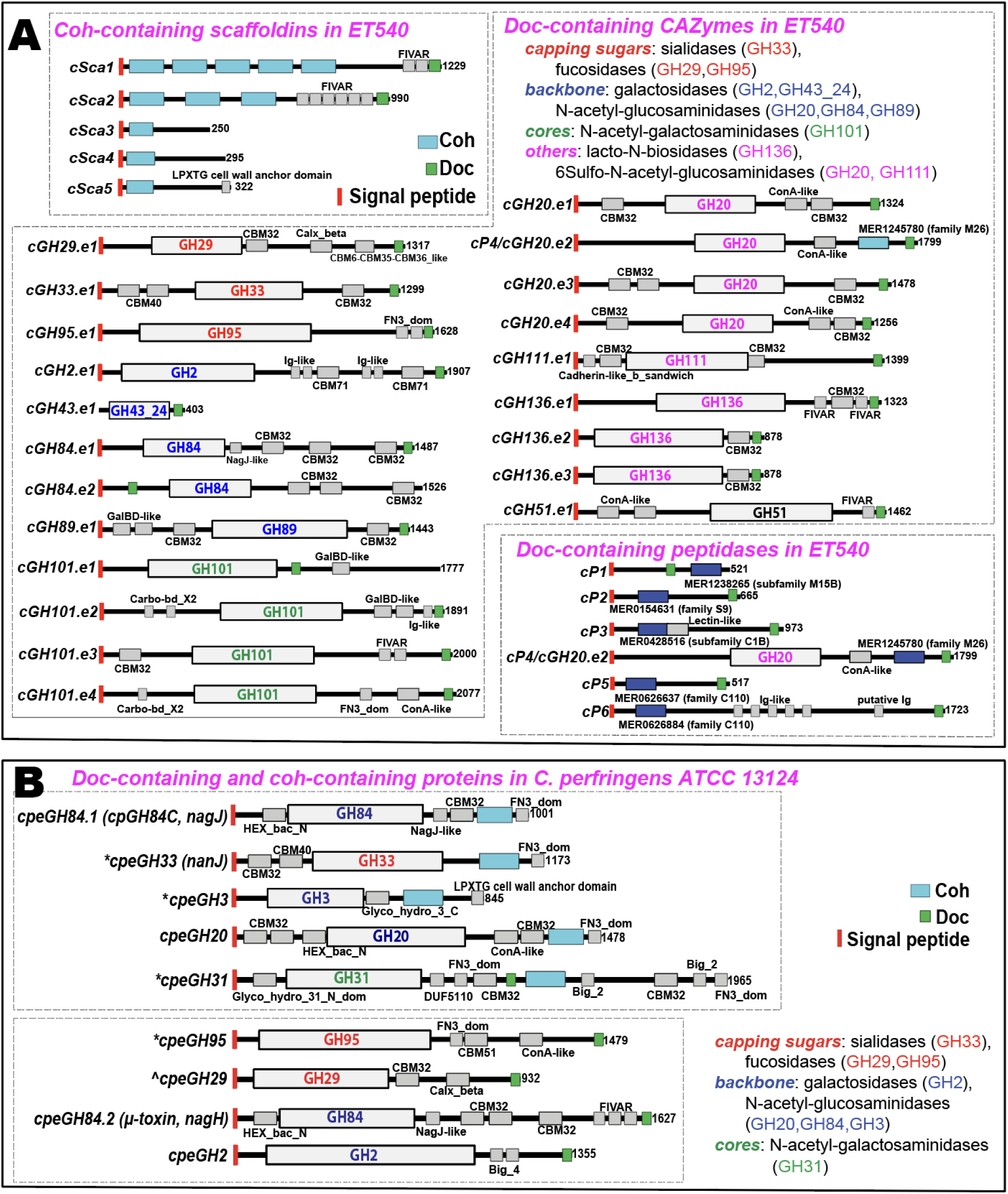
Comparison of domain diagrams of Doc-containing and Coh-containing proteins in two genomes. (**A**) ET540 and (**B**) ATCC 13124. For Doc-containing proteins, only those with CAZyme and peptidase domains are shown. CAZyme domains are colored based on their biochemical activities from literature (2, 23, 24) and CAZy website (cazy.org) (25). * indicates that these proteins have a weak cohesin or dockerin signal that failed to be identified by the Pfam cohesin and dockerin HMMs with E-value < 1e-4 and coverage > 60% (Figure 1A). ^ indicates the GH29 protein that failed to be identified with the dockerin domain in the previous study (39).

*Clostridium perfringens* ATCC 13124 has been previously shown to use Coh-Doc interactions to form multi-toxin complexes for animal tissue destruction (*39*). In this context, ATCC 13124 has five Coh-containing proteins, each having a single Coh module, previously known as X82 (*39*). Interestingly, all the five Coh-containing proteins also have a GH domain (**Figure 3B**), while no GH domains are present in Sca proteins of ET540 (**Figure 3A**). Therefore, these Coh-containing proteins in ATCC 13124 are very different from traditional scaffoldins, which usually contain no catalytic domains but only multiple Coh tandem repeats. The only reported exception is scaffoldin CipV from *Acetivibrio cellulolyticus* (*40*) that contains a GH9 domain in addition to seven Coh repeats. ATCC 13124 also has five Doc-containing GHs, including one GH29 protein that was overlooked in the previous study (*39*) and one GH31 protein with both Coh and Doh domains. The five GH proteins include two known toxins in *C. perfringens*: a hyaluronidase (μ-toxin, NagH, GH84) and a sialidase (NanJ, GH33) (*41*). Overall, ATCC 13124 has a very distinct and much simpler Coh-Doc interaction system (**Figure 3B**) than ET540.

ET540’s mucinolysome system has a total of 12 mucin glycan-degrading GH families (**Figure 3**, also indicated by “*” in **Figure 2C**). Eight other GH families were also reported in the well-studied mucin glycan-degrading gut bacteria *A. muciniphila*, *C. perfringens*, *Bifidobacterium bifidum*, *R. gnavus,* and *Roseburia inulinivorans* (*2, 6, 23, 24*). These eight GH families were found in *Limousia* CAZymes without Doc domains. For example, GH42 (β-galactosidases) and GH112 (lacto-N-biose phosphorylases) were present in most of the 65 *Limousia* genomes. Some families (e.g., GH3 β-N-acetylhexosaminidases and GH123 β-N-acetylgalactosaminidases) were almost exclusively found in one of the three *Limousia* species (sp944380855, **Figure 2C**), with other families (GH31 and GH36 α-N-acetylgalactosaminidases, GH95 and GH51 fucosidases) almost exclusively absent in sp944380855. Overall, GH2, GH20, GH84, GH101, and GH136 tend to have multiple protein copies in each *Limousia* genome, while other families (e.g., GH29, GH33, GH42, GH43, GH89, GH31, GH112, GH51) have only one copy in each genome (**Figure 2C**).

We further compared the total CAZyme repertories of the two representative *L. pullorum* genomes (MGYG000005496, or *Lpuh,* from human, and ET540, or *Lpuc,* from chicken) against those of *A. muciniphila*, *C. perfringens* and *R. gnavus* (**Figure 2D** and **Table S3**). The two *Limousia* genomes encode 23 and 24 GH families for degradation of glycans from different sources (34 and 45 GH proteins in *Lpuh* and *Lpuc*, respectively), lower than the numbers of GH families found in the three known mucin glycan-degrading genomes (*A. muciniphila*, *C. perfringens*, *R. gnavus*). However, when only considering the percentage of mucin glycan-degrading GHs (all rows in **Figure 2C** and all colored GH families in **Figure 2D**) over all genes in the genome, *Limousia* genomes have similar percentages (1.1% and 1.3%) as *C. perfringens* (1.2%), higher than *R. gnavus* (0.7%), but lower than *A. muciniphila* (2.4%). The percentage of mucin glycan-degrading GHs over all GHs is also highest in *A. muciniphila* (83.1%), followed by *Lpuc* (68.9%). Extracellular activities of GHs are important for mucin degradation (*42*). We found that *A. muciniphila* has the highest percentages of GHs (68.85%) and mucin glycan GHs (78.43%) predicted to be secreted, followed by *Lpuc* (48.89% and 70.97%) (**Figure 2D, Table S3**). Overall, 95.24% and 100% secreted GHs in *A. muciniphila* and *Lpuc* are predicted to target mucins.

*A. muciniphila* has more copies of GH29, GH2, GH20, and GH16, and more unique families (GH181, GH27, GH35, GH97, GH110, and GH109) (**Figure 2D**). *Limousia* genomes also have unique families absent in the other genomes, such as GH136 and GH111. *Limousia* and *C. perfringens* are from the *Eubacteriales* order, and share GH101, absent in the other two genomes, which are from the *Lachnospirales* order of Firmicutes (*R. gnavus*) and from the Verrucomicrobia phylum (*A. muciniphila*). *Limousia* and *C. perfringens* genomes also more CBM32, but less GT2 and GT4 proteins than the other genomes. All five genomes share GH13 and GH77 for α-glucan degradation (**Figure 2D**). *Limousia* genomes encode no PLs (polysaccharide lysases) and AAs (CAZyme of auxiliary activities), and very few CEs (carbohydrate esterases).

### AlphaFold3 predicts Coh-Doc interactions in the mucinolysome system of ET540

To better understand the potential for mucinolysome components to interact, we predicted the 3D structures for all cohesin and dockerin modules and their interactions in the mucinolysomes of *L. pullorum* ET540 and *C. perfringens* ATCC 13124 using AlphaFold3 (AF3) (*43*). The predicted structures of Coh-Doc complexes were further processed by FoldDock (*44*) to calculate pDockQ (predicted docking model quality) scores for all Coh-Doc pairs. The pDockQ score measures the confidence of PPI (protein-protein interaction) by considering the PPI interface pLDDT (predicted Local Distance Difference Test) score (IF_pLDDT) and the number of residues in the PPI interface (IF_contacts). Previous reports showed that PPI models with pDockQ > 0.23 are 70% likely correct (*45*) and considered as acceptable (*44*).

We first tested the feasibility of this approach on ATCC 13124 as a proof of concept. Previously, Coh-Doc interactions have been experimentally detected in ATCC 13124 (*39*), showing that the Coh-containing GH84.1 (NagJ), GH33 (NanJ), and GH20 proteins possessed strong binding properties to the Doc-containing GH84.2 (NagH), GH95, and GH2. The other two Coh-containing proteins (GH31 and GH3) exhibited no binding to Doc-containing proteins. With a pDockQ > 0.3 threshold, AF3 predicted all these experimentally validated PPIs in ATCC 13124 (**Figure 4A**). Additionally, AF3 predicted PPIs between GH31-Coh and two dockerins: GH31-Doc, and GH2-Doc. The interfaces of these two PPIs are ∼18% disordered (**Table S4**), and therefore the PPIs might indeed exist but failed to be detected using an affinity ELISA-based assay (*39*). For the newly identified GH29-doc, AF3 predicted interactions with GH84.1 (NagJ)- Coh, GH33 (NanJ)-Coh and GH20-Coh.

**Figure 4:**
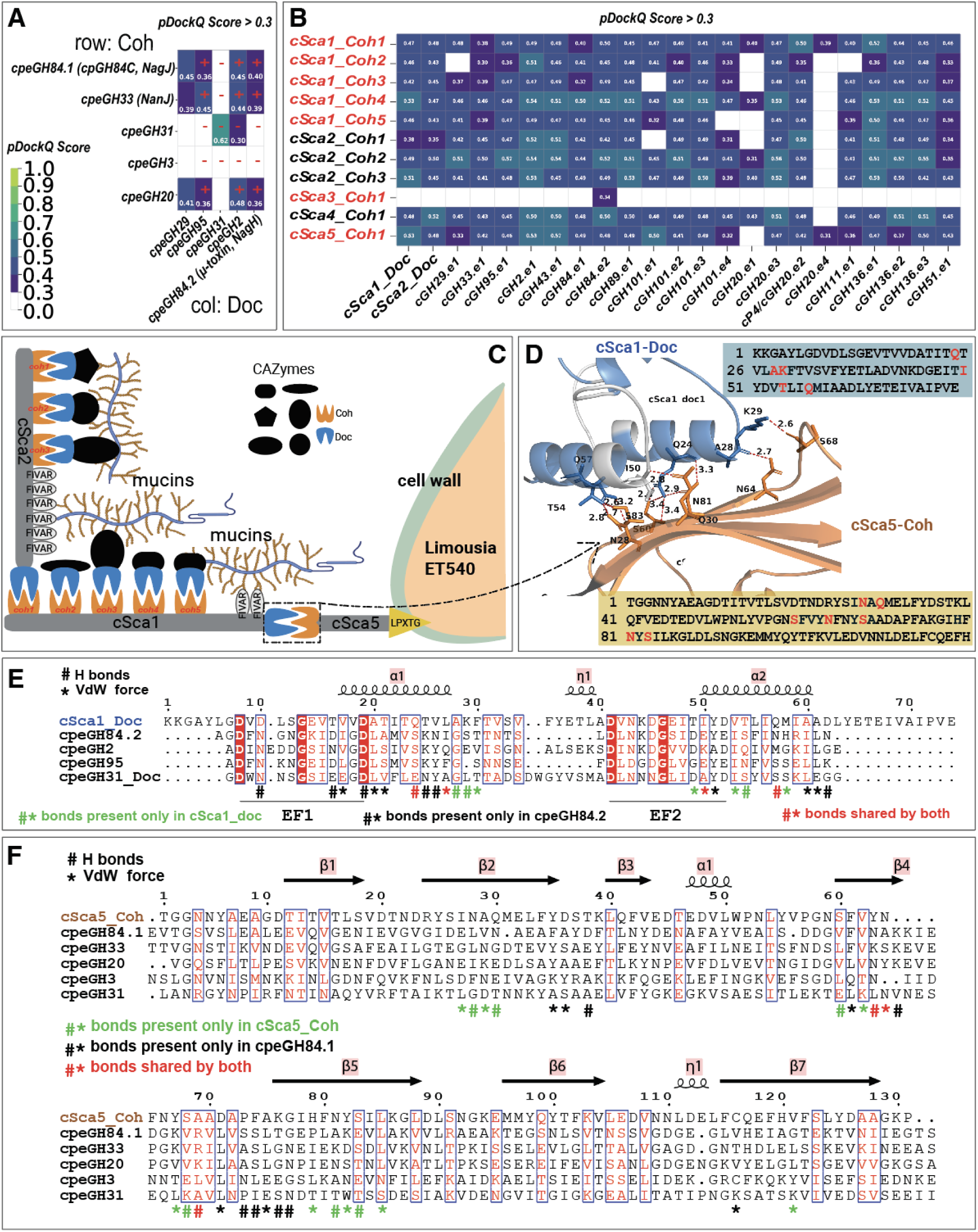
AlphaFold3 predicted functional Coh-Doc interactions. Dockerin and cohesin domain sequence pairs were used as input to AlphaFold3 for 3D structure and interaction predictions. The resulting PDB files were input to FoldDock to calculate the pDockQ scores. Scores larger than 0.3 are shown. (**A**) Protein-protein interactions (PPIs) are predicted between cohesins (rows) and dockerins (columns) in GHs of C. perfringens ATCC 13124. “+” and “-“ indicate the presence and absence of experimentally characterized PPIs (39). (**B**) PPIs are predicted between cohesins in scaffoldins (rows) and dockerins in GHs (columns) of ET540. (**C**) A conceptual model is proposed illustrating the protein organization in the mucinolysome of ET540. (**D**) Predicted PPI interface between cSca1-Doc and cSca5-Coh. Hydrogen-bond contacts are shown as red dashed lines with corresponding distance < 3.5 Å between atoms. Sequences of cSca1-Doc and cSca5-Coh are shown beside the PPI interface structure with residues highlighted corresponding to residues in the hydrogen-bond contact interface. (**E**) Sequence alignment of cSca1-Doc against dockerin sequences in four ATCC 13124 proteins. Key residues involved in hydrogen-bonding (#) and van der Waals (*) contacts are indicated. Residues for cSca1-Doc are predicted, while residues for ATCC 13124 proteins were published previously (39). The two EF hand motifs are indicated. (**F**) Sequence alignment of cSca5-Coh against cohesin sequences in five published ATCC 13124 proteins (39).

As all experimentally validated Coh-Doc PPIs in ATCC 13124 were predicted by AF3 with the threshold pDockQ > 0.3, we applied this threshold to AF3 predicted PPIs (**Table S5**) between dockerins and cohesins in ET540 (**Figure 4B**). Most Coh-Doc pairs were predicted to have interactions supported by higher pDockQ scores (compared to those in ATCC 13124), except for cSca3-Coh and cGH20.e4. Notably, cSca5 has a C-terminal LPxTG cell wall sortase anchoring domain, which is key to covalently link the protein to peptidoglycans of bacterial cell walls (*46*). Interestingly, the N-terminal cSca5-Coh was predicted to interact with both cSca1-Doc (pDockQ=0.53, IF_pLDDT=96.4, IF_contacts=57) and cSca2-Doc (pDockQ=0.48, IF_pLDDT=94.7, IF_contacts=53), as well as dockerins of most GHs. cSca1 and cSca2 both have multiple cohesin modules, predicted to interact with their own dockerins (i.e., cSca1-doc and cSca2-Doc) and those of most of the GHs. Thus, through Coh-Doc PPIs among these cSca proteins and GHs, multi-protein mucinolysome complexes of varying components could form and be anchored onto bacterial cell walls through cSca5 (**Figure 4C**).

We further compared the predicted PPI interface of cSca5-Coh and cSca1-Doc with the experimentally determined PPI interface of cpeGH84.1-Coh and cpeGH84.2-Doc (*39*). Key residues in the two EF hand motifs (**Figure 4D**) were found to be conserved between ATCC 13124 and ET540, while those involved in the Coh-Doc interactions were not very conserved but clustered in the sequence alignments (**Figure 4E,F**). This may reflect the species specificity in terms of Coh-Doc interactions to form the multi-protein complex.

### ET540 grows with mucin as the sole carbon source

To investigate mucinolysome function *in vivo*, the ability of strain ET540 to grow with mucin as the sole carbon source was initially evaluated using defined basal medium (DBM, **Table S6**) on agar plates without an added carbon source or with 0.2% w/v porcine gastric mucin (**Figure S2**). From these experiments, we observed that mucin enhanced the growth of ET540 by more than 10^5^-fold, with no colonies recovered on DBM without an added carbon source (<10 CFU/mL).We further measured the growth of ET540 in DBM broth with different concentrations of mucin over time (**Figure 5A**). We observed a linear increase in maximal cell density (measured as OD_625_) as mucin concentrations increased (**Figure 5B**); similar results were observed in an independent experiment (**Figure S3**). We also observed modest decreases in doubling time during the exponential phase with increasing mucin concentrations (**Figure 5C**).

**Figure 5:**
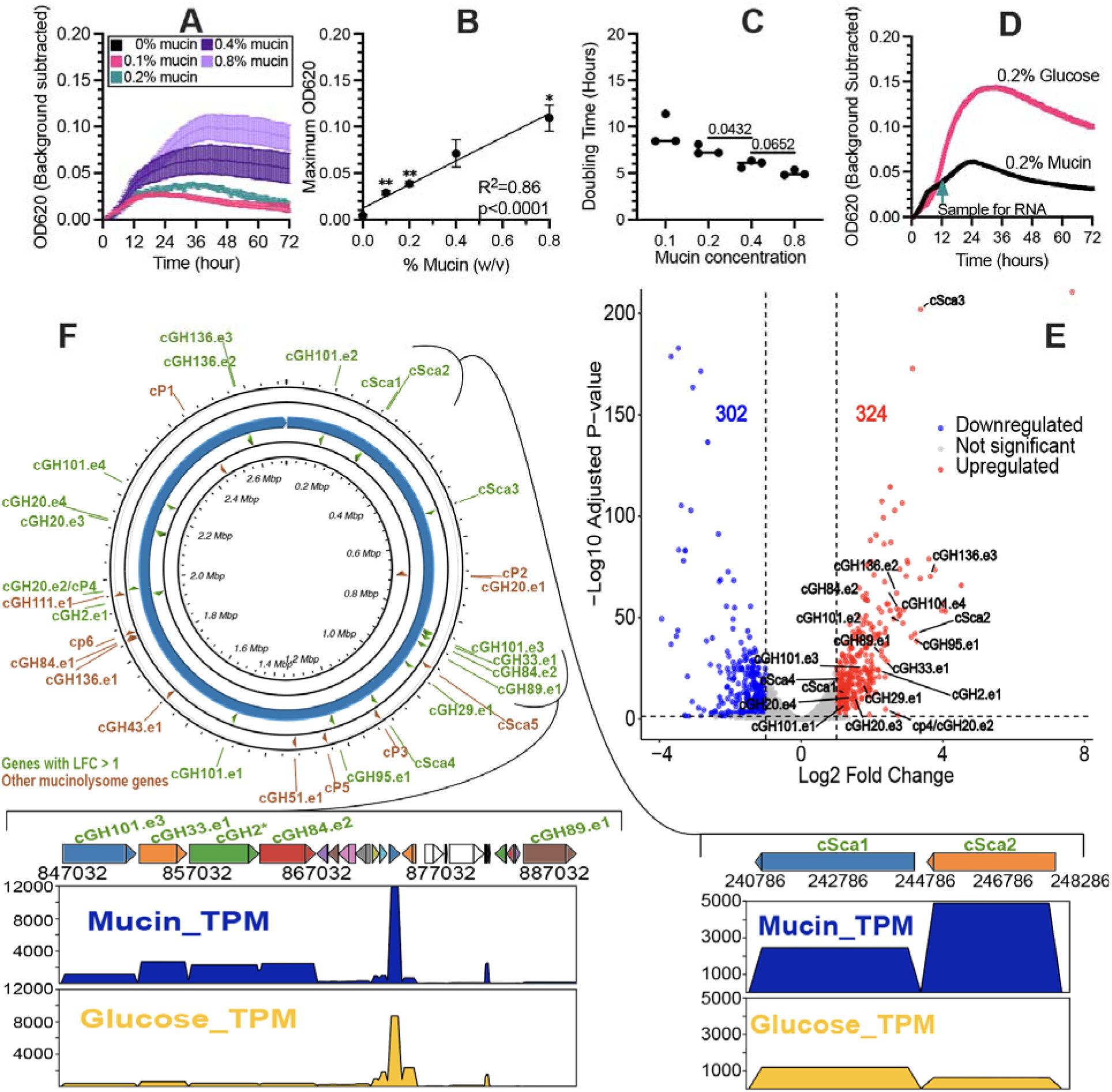
ET540 grows with mucin as the sole carbon source with most mucinolysome genes up-regulated. In (**A**)-(**C**), ET540 was grown in DBM + 0.2% mucin for 48 hr, diluted into fresh DBM medium containing the indicated mucin concentrations, where (**A**) growth over time in culture, (**B**) maximum OD_620_ relative to mucin concentration, and (**C**) doubling time from 2-24 hr for each mucin concentration were plotted. In (**A**), the mean ± SEM values are plotted for triplicate cultures at each time point. In (**B**), mean ± SEM values are plotted for triplicate cultures at each mucin concentration. Linear regression was used to determine the goodness of fit (R^2^) and significance of slope deviation from zero, with p<0.05 reported. Significance of differences in maximum OD_620_ values at each mucin concentration relative to no mucin was determined by one-way ANOVA with Brown-Forsythe correction for unequal variances and Dunnett’s T3 correction for multiple comparisons, with ** indicating p<0.01. In (**C**), each point represents a replicate with the line indicating the median. Significance of difference in doubling time between mucin concentrations was determined by one-way ANOVA with Brown-Forsythe correction for unequal variances and Dunnett’s T3 correction for multiple comparisons, with p values <0.1 reported. In (**D**), growth over time in culture was plotted as the mean ± SEM and the time of sample collection for RNA extraction is indicated by an arrow. (**E**) Differentially expressed genes (DEGs) between growth on glucose (DBM #2 + 0.2% glucose) and on mucin (DBM #2 + 0.2% mucin) are shown in the volcano plot. The horizontal line indicates the adjusted p-value (FDR) threshold of < 0.05. Vertical lines indicate the log2-fold change log2FC > 1 (two times higher in mucin, right line) or log2FC < −1 (two times lower in mucin, left line). Upregulated mucinolysome genes under growth on mucin are indicated (**Table 2**). (**F**) Circular view of genomic locations of all mucinolysome CAZyme and peptidase genes. Those with log2FC > 1 are shown in green. Read mapping coverage plots are shown for two gene clusters with upregulated mucinolysome genes. The y-axis shows the TPM values and the x-axis shows the positions in the genome.

**Table 2:**
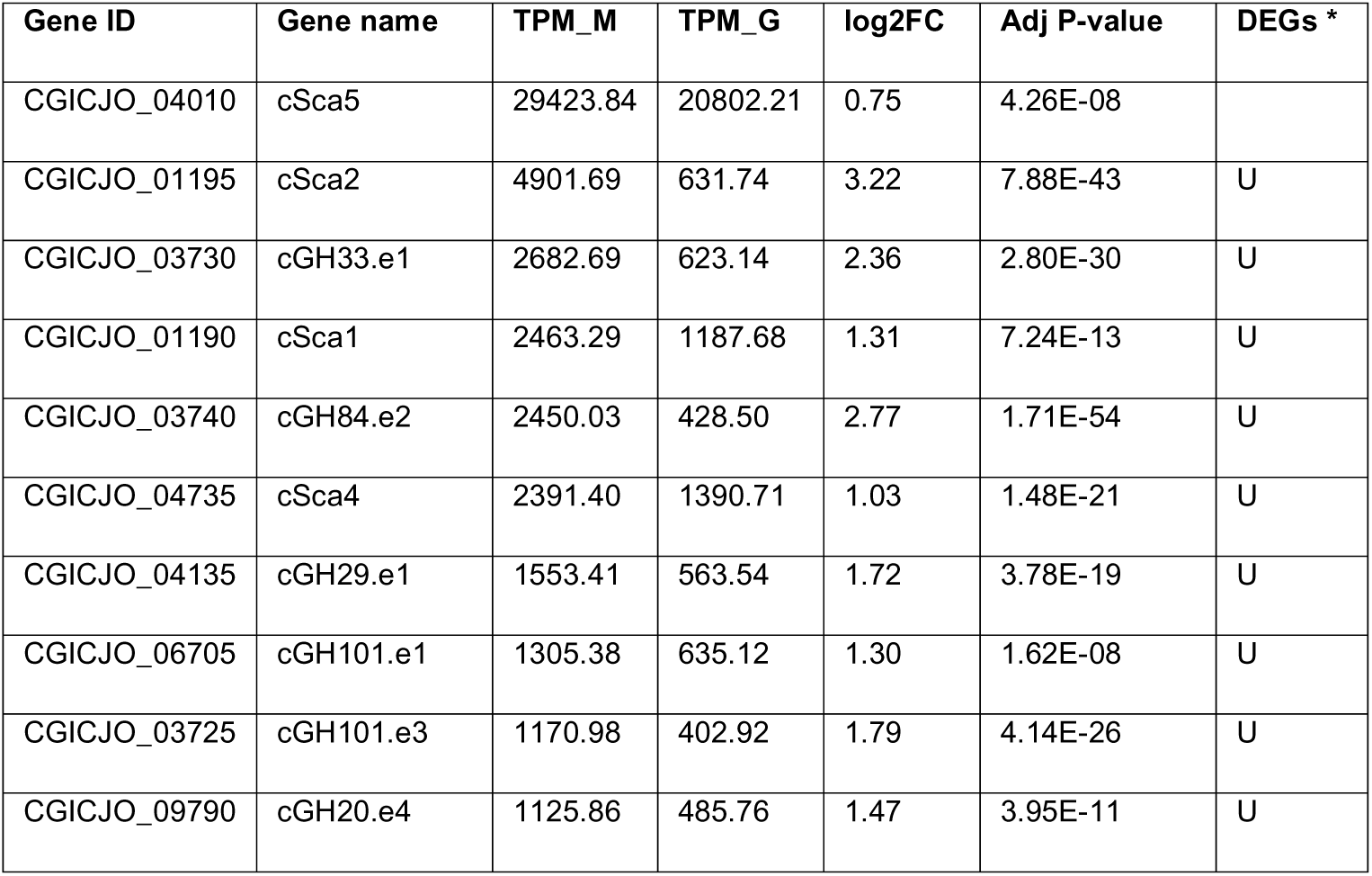
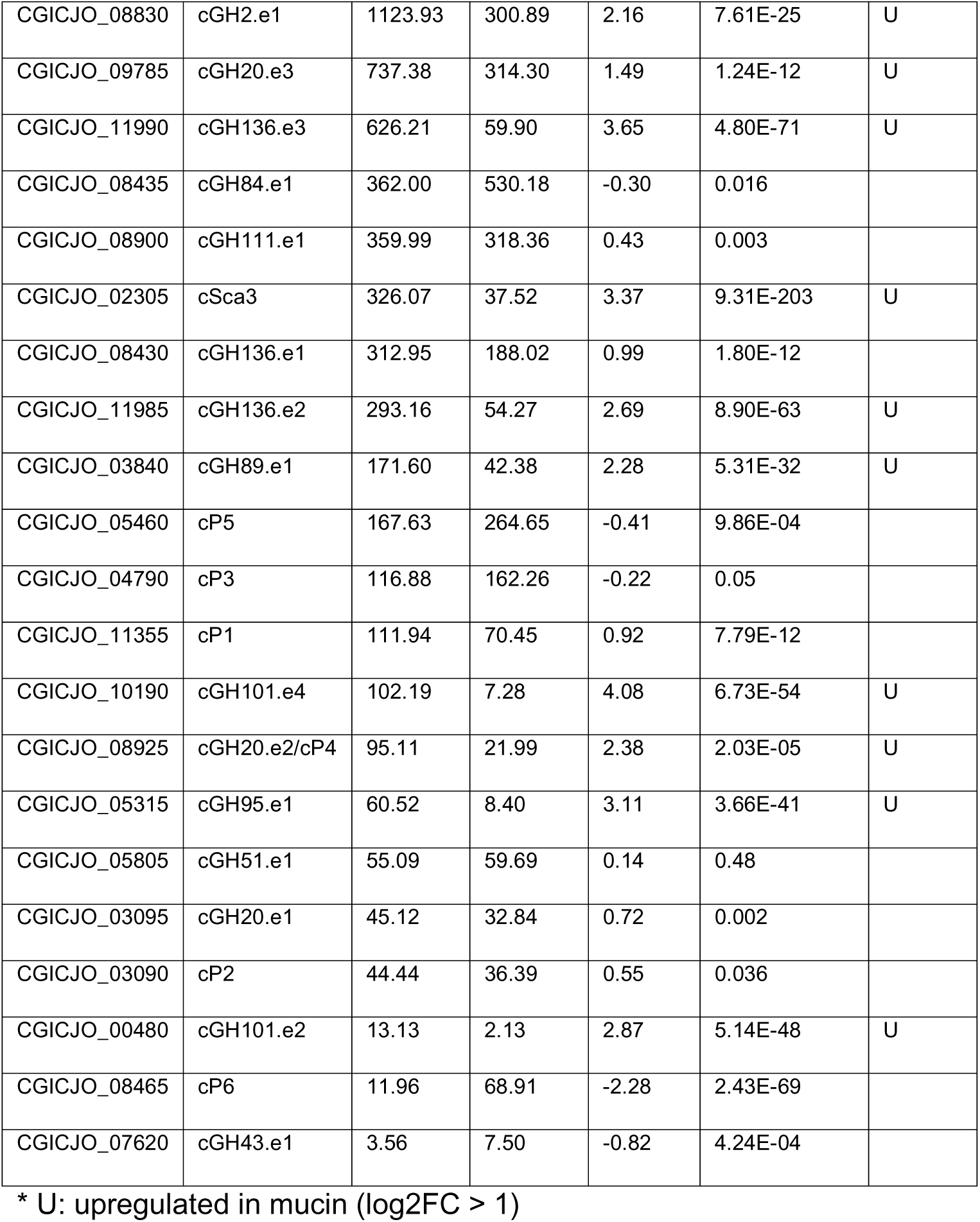
Mucinolysome gene expression in ET540 sorted by TPM_M (transcripts per kilobase million when grown on mucin).

Growing ET540 in DBM with added glucose, we observed that concentrations as high as 5% glucose did not support growth (**Figure S4A**). Similar observations have been reported for *A. muciniphila*, where it was observed that supplementation with 0.2% tryptone would allow cells to metabolize glucose (*47*). We evaluated growth with glucose as a carbon source in DBM supplemented with 0.2% tryptone (DBM #2, **Table S6**) and observed glucose-dependent growth at 0.25%, 0.5%, and 1% glucose (**Figure S4B**), with similar maximum cell densities (**Figure S4C**) and doubling times (**Figure S4C**) at different concentrations.

To optimize growth conditions for comparing gene expression with mucin or glucose as the primary carbon source, we compared ET540 growth in DBM #2 with 0.1% or 0.2% mucin or glucose (**Figure S5A**) as ET540 did not grow in DBM with glucose. While we observed higher growth yields (**Figure S5B**) and shorter doubling times (**Figure S5C**) for cells grown in glucose compared to mucin, although we observed that cell densities were similar between ET540 in 0.2% mucin or 0.2% glucose after 12 hr of growth (**Figure 5D**). We used these conditions for growth of strains for RNA extraction and sequencing (described below) and collected growth measurements of replicate cultures grown in parallel (**Figure S5D-5F**), with data from this replicate experiment exhibiting similar patterns of growth.

### Most mucinolysome genes are upregulated in ET540 grown on mucins

With the two growth conditions indicated in **Figure 5D**, we generated 11∼14Gb short-read RNA-seq data per sample and mapped clean reads to the ET540 genome. Using DESeq2 (*48*), 626 (out of 2,527) genes (**Table S7**) were identified as differentially expressed genes (DEGs) between the glucose group and the mucin group using an adjusted p-value (FDR) threshold of < 0.05 and an absolute log2-fold change ∣log2FC∣ > 1 (2-fold change). There were 324 upregulated DEGs and 302 downregulated DEGs in mucin compared to glucose (**Figure 5E**). Enriched KEGG functions found in the upregulated DEGs included those associated with “Glycosaminoglycan degradation”, “Galactose metabolism” and “Other glycan degradation”, which are all related to mucin degradation.

Interestingly, 19 (71.1%) of the 26 scaffoldin + Doc-containing CAZyme genes were upregulated DEGs in the presence of mucin (**Table 2**). Four of the five scaffoldin-encoding genes are DEGs (**Figure 5E**). The only exception is cSca5, which indeed has a higher expression in mucin samples (FC=1.69), but does not meet the log2FC>1 threshold (log2FC=0.75, **Table 2**). Most interestingly, the cSca5-encoding gene has the highest expression among all the 2,527 ET540 genes (**Table S7**) with a TPM (Transcripts Per kilobase Million) value = 29,423 in mucin samples; the cSca5-encoding gene also has the second highest expression among all genes with a TPM = 20,802 in glucose samples. The cSca5-encoding gene has an expression six to seven times higher than the second highest expressed mucinolysome gene (cSca2 in mucin samples and cSca4 in glucose samples, respectively, **Table 2**). This very high and constitutive gene expression of cSca5 (with a cell wall sortase anchoring motif) in both conditions strengthens its critical role in anchoring scaffoldin complexes to the bacterial cell wall and promoting the assembly of the mucinolysome and other potential degradative complexes (i.e., peptidases and other protein domains, **Table S2**).

Additionally, genes encoding six Doc-containing CAZymes (cGH84.e1, cGH111.e1, cGH136.e1, cGH20.e1, cGH43.e1, cGH51.e1) and five Doc-containing peptidases (cP1, cP2, cP3, cP5, cP6) were not upregulated DEGs (**Table 2**), but like the gene encoding cSca5, many of these genes had higher expression in mucin samples but did not meet the log2FC>1 threshold.Interestingly, many mucinolysome genes form physically linked and co-expressed gene clusters (**Figure 5F**). For example, the genes encoding cSca1 and cSca2 are 287 base pairs apart and are co-upregulated in mucins. Three mucinolysome genes, encoding cGH101.e3, cGH33.e1, and GH84.e2 form a gene cluster with a non-mucinolysome GH2 gene, with all four genes upregulated upon growth on mucins (**Table S7**). The gene encoding cGH89.e1 is located 17 genes downstream of this cluster, also upregulated upon growth on mucins (**Table S7**). Overall, the RNA-seq experiment demonstrated that most mucinolysome genes were highly expressed and upregulated when ET540 was grown on mucins.

### Prevalence and abundance of mucinolysome-encoding *Limousia*

To study the abundance of these mucinolysome-encoding *Limousia* MAGs, we downloaded the original metagenomic reads used to build these MAGs (if available) and mapped reads back to these MAGs (see **Methods**). We found that, for the seven human MAGs, their abundances were low (0.06%-0.61%) in their original samples (**Figure 6A**, **Table 1**). For MAGs from other animals, their abundances range between 0.0098% and 12.51% (**Figure 6A**). MAGs from cows had higher abundances, followed by MAGs sourced from pigs and humans. The chicken MAGs had the lowest abundances in their original samples. Despite the low abundance, most MAGs had almost 100% mapping breadth of coverage (**Figure 6B**), verifying their existence in the original samples. Overall, the mucinolysome-encoding *Limousia* bacteria have low abundance in different animal hosts (<1% for most MAGs).

**Figure 6:**
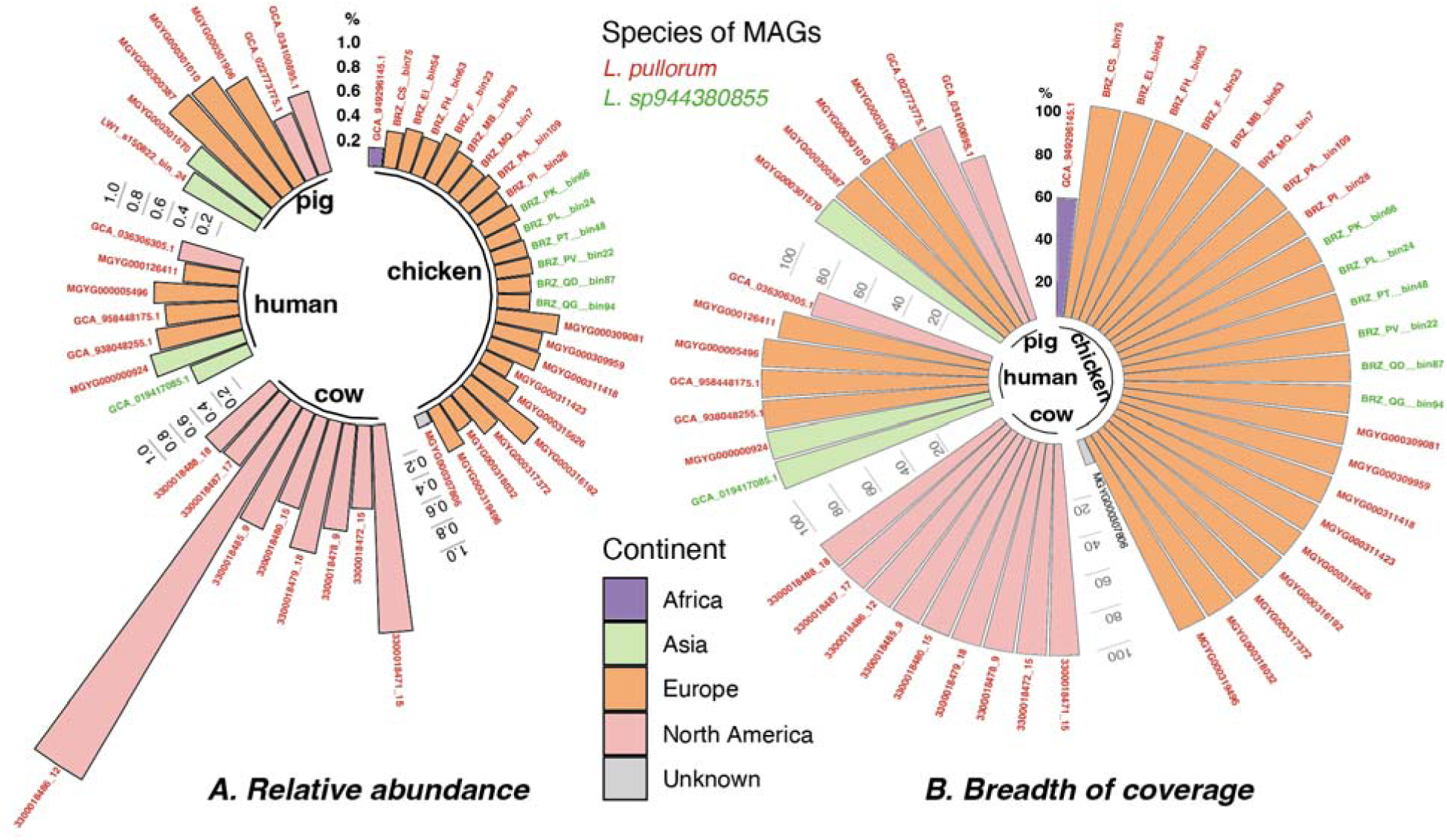
Relative abundance of Limousia MAGs by mapping reads of original samples. For each sample, all clean reads are mapped to the respective MAG using Bowtie (see Method). The mapped reads are processed to calculate the relative abundance (RA) and the breadth of coverage (BC) for the MAG in the sample. (**A**) RA of a MAG is calculated as the count of the mapped reads to the MAG divided by the count of all reads in the sample. (**B**) BC of a MAG is calculated as the count of bases that have at least one read mapped divided by the count of all bases in the MAG (i.e., the length of the MAG). The bars indicate the MAGs and are colored according to source continents. Bar labels are the MAG names that are colored based on species defined in Figure 2A. Original read samples for 13 chicken MAGs failed to be located and were excluded in this analysis. Three MAGs (GCA_034100895.1, GCA_036306305.1 and MGYG000126411) that were assembled from multiple samples had their abundances in each of the samples calculated and then averaged.

The prevalence of mucinolysome-encoding *Limousia* were studied by mapping metagenomic shotgun reads of 2,897 fecal samples from various human cohorts and wild/domesticated animals (**Figure 7A**). To reduce the computational cost, we subsampled 10 million reads from each sample and mapped the clean reads to three different reference datasets (**Figure 7B**): (i) the entire genomes of 65 *Limousia* MAGs, (ii) 16S RNA sequences of four MAGs (no 16S RNAs were found in other MAGs), and (iii) coding sequences (CDSs) of five scaffoldin proteins and 21 GHs of ET540. Using inStrain (*49*) and CoverM, we calculated the breadth of coverage (BC) and average read depth (ARD) for each MAG, 16S RNA, and CDS in each sample (see **Methods**).

**Figure 7:**
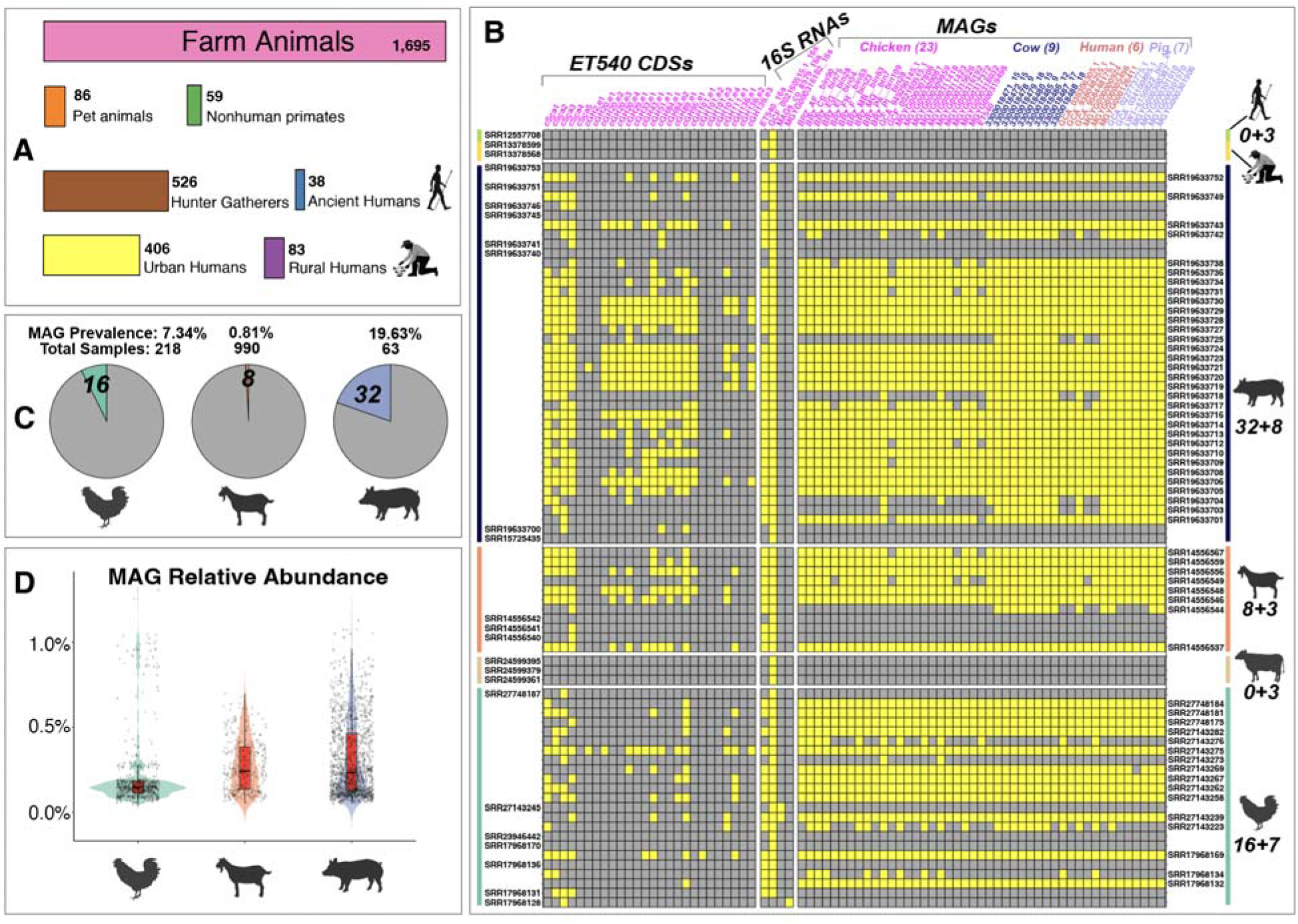
Prevalence and abundance of Limousia MAGs in different human and animal hosts. (**A**) The total fecal sample count is 2,897. Sample counts from different animal and human hosts are shown as bars. (**B**) Heatmap of read mapping against MAGs, 16S RNAs, and ET540 mucinolysome CDSs as reference sequences. Yellow color means mapped or positive mapping and gray color means unmapped. Rows are sample SRA accessions, and columns are references (MAGs/16S RNAs/CDSs). Rows are ordered by hosts. SRAs shown on the right side are samples mapped to MAGs, and SRAs shown on the left side are samples unmapped to MAGs but mapped to 16S RNAs or CDSs. Two numbers are shown on the right side (e.g., 32+8): the first number is the count of samples (SRA IDs on the right) mapped to MAGs, and the second number is the count of samples (SRA IDs on the left) unmapped to MAGs but mapped to 16S RNAs or CDSs. Reference IDs are color-coded by host origin. (**C**) MAG prevalence (positive sample count divided by total sample count) in different hosts is shown as pie charts. (**D**) Boxplot of MAG relative abundance in positive samples grouped by hosts.

As a result of the above-described approach, a total of 56 samples were found to be mapped to a total of 45 MAGs (**Figure 7B**). These 56 samples were therefore considered to encode the mucinolysome system. Note that cross-mapping could occur, meaning read samples can be mapped to MAGs of different hosts. For example, the eight sheep samples were mapped to MAGs of chicken, cow, human, and pig meeting our thresholds (BC of ≥0.6 and ARD of ≥1). The prevalence of mucinolysome in different host samples was calculated as the number of positive samples divided by the total number of samples (**Figure 7C**). We observed the presence of MAGs in 16 chicken samples (prevalence rate = 7.34%), eight (0.81%) goat samples, and 32 (19.63%) pig samples. No samples from cow, human, non-human primate, and other animal samples were positive. This was unexpected since our 65 *Limousia* MAGs contained seven MAGs from human and eight MAGs from cow (**Figure 2A**). Overall, a total prevalence rate of 1.94% was observed across all samples. All the 56 positive samples were sourced from China except for one from Germany. Notably, seven out of the 32 pig samples (**Figure 7C**) were derived from animals reported to have diarrhea and the rest were from healthy piglets, but no disease information was found available for chicken and goat samples.

In addition to MAGs, we also mapped clean reads to 16S RNA sequences of four MAGs and ET540 CDSs of five scaffoldin proteins and 21 GHs. All the positive 56 samples mapped to the MAGs were confirmed by positive mapping to at least one 16S RNA and one CDS (**Figure 7B**). Interestingly, we also found 24 new samples mapped to at least one 16S RNA or one CDS. These 24 samples included three human samples and three cow samples, which were only mapped to 16S RNAs, meeting the stringent thresholds: BC of ≥0.6, ARD of ≥1, and read alignment sequence identity > 99%. Given the low abundance of mucinolysome-encoding *Limousia*, it is likely that these are true positive samples for the presence of mucinolysomes. The presence of mucinolysomes in pig, chicken, and goat was well supported despite the low prevalence (**Figure 7C**) and abundance (**Figure 7D**) in read mapping.

## DISCUSSION

Mucins form a protective layer covering the epithelial surfaces in humans and animals. In the GI tract, mucin glycans have been shown to be degraded by mutualistic microbes, such as *A. muciniphila* and *B. thetaiotamicron*, and pathogens, such as *C. perfringens*. Mucin glycan degradation by pathogens can contribute to different disease states, and thus the discovery of a new molecular mechanism for mucin glycan degradation has a potential major impact in biomedicine, particularly for understanding the balance between mucin glycan degradation that enhances barrier function and degradation that increases susceptibility to disease. In this study, we uncovered a new potential molecular mechanism that *Limousia* bacteria, present in human and animal GI tracts, employ for mucin glycan degradation. In this mechanism, *Limousia* spp. encode mucinolysomes, which are large extracellular multi-enzyme complexes formed through hallmark Coh-Doc interactions for processive degradation of mucin by mucin glycan-degrading CAZymes and peptidases (**Figure 4C**). This contrasts with other well studied gut bacteria, such as *Bacteroides* spp., which use the polysaccharide utilization loci (PUL) mechanism for mucin utilization, where genes encoding CAZymes, transporters, and regulators form co-regulated gene clusters in genomes (*7, 8*). Mucin utilization loci (MUL) were also recently discovered in the most studied mucin-degrading bacterial species, *A. muciniphila* (*42*). However, these MULs do not encode CAZymes. Instead, they encode type IV-like pili proteins and transporters, which might be key to transporting mucinolytic enzymes to the outside or to the periplasm of the cell, and to importing mucin glycans and their partially degraded products into the cell to form the intracellular compartment called mucinosomes for further degradation (*42*).

The mucinolysome mechanism discovered in *Limousia* bacteria follows the classical cellulosome paradigm well described in *Clostridium* and *Ruminococcus* bacteria (*14*). Our study presents the first evidence that the cellulosome architecture is not only used for the degradation of plant polysaccharides (lignocelluloses and starches) but can also be utilized in animal glycan mucin degradation. Although Adams *et al.* previously showed that *C. perfringens* ATCC 13124 uses Coh-Doc interactions to form complexes for degradation of host-associated glycans (*39*), these complexes (**Figure 3B**) do not appear to follow the classical cellulosome architecture with assembly of multiple Doc-containing CAZymes on scaffoldin proteins with multiple Coh-domains. Rather, this organization appears to facilitate pairwise interactions between GH proteins. In contrast, in *Limousia* ET540, the classical cellulosome architecture is preserved, as the five scaffoldins do not have GH domains, and tandem Coh repeats are present in cSca1 and cSca2.

Structure-based prediction of Coh-Doc interactions, validated by known interactions in *C. perfringens* ATCC 13124 (*39*), was further applied to predict Coh-Doc interactions in *Limousia* ET540. The results revealed that most of the scaffoldins (except cSca3) could have interactions with all mucin degrading CAZymes and peptidases (**Figure 4B**). It is interesting that the longest cSca1 and cSca2, which contain five and three cohesin repeats, respectively, can potentially use their C-terminal dockerins to interact with the cell wall-anchoring cSca5. These interactions will likely play a key role in the assembly of the large multi-scaffoldin mucinolysome system and its attachment to the bacterial cell walls. More interestingly, the gene for the cell wall-anchoring cSca5 was the most highly expressed gene among all genes in the genome (**Table 2**, **Table S7**, ranking 1^st^ in mucin- and 2^nd^ in glucose growth media), and was upregulated (1.7-fold) in mucin-containing growth media. The genes for the other four scaffoldins were all significantly upregulated (>2-fold) in mucin-containing growth media. While future experimental validations of these interactions are needed, our initial mucinolysome model (**Figure 4C**) likely underestimates the complexity of the real mucinolysome system in Nature.

In addition to CAZymes, peptidases and sulfatases are also important for complete mucin degradation. The *Limousia* ET540 genome has a total of 92 peptidases according to a BLASTP search against the MEROPS database (E-value < 1e-10), and nine sulfatases according to a hmmsearch against the SulfAtlas database (E-value < 1e-3) (**Table S7**). Six of the peptidases contain dockerins (**Figure 3A**), and six peptidases were upregulated in ET540 grown in the presence of mucin. No sulfatases contain dockerins, and none were upregulated in upon growth on mucin. It is known that very few bacterial sulfatases have been experimentally characterized and are thus less represented in the databases (*50*). Given that the ET540 isolate can grow well on mucins, novel sulfatases that do not share sequence similarity with known enzymes are likely to be discovered in *Limousia*.

This study characterizes and adds the *Limousia* bacteria as a unique member to the list of human gut mucin glycan-degrading bacteria (*23*). Its genome encodes the classical cellulosome-like system for mucin glycan degradation (i.e., mucinolysome) not found in other mucin glycan-degrading bacteria. Other mucin degraders, such as *A. muciniphila,* exhibit a single dockerin and a single cohesin encoded in its genome, within a GH31 protein (Amuc_1008/WP_012420049) (*6*), whereas *R. gnavus* lacks any dockerins or cohesins. As described above, mucin-glycan degrading Coh and Doc domain proteins are present in *C. perfringens*, but these proteins do not appear to form a mucinolysome. Compared to *C. perfringens* and *R. gnavus, Limousia* ET540 has higher percentages of mucin glycan-degrading GHs and secreted GHs (**Figure 2D, Table S3**), while still being lower than those of *A. muciniphila*, one of the most well characterized mucin glycan-degrading bacteria. Future experimental studies are needed to compare the competitive growth of these different mucin glycan degraders. *Limousia* represents a new model microorganism to study the highly organized mucin degradation using the novel mucinolysome mechanism.

Our comprehensive read mapping-based search found that the mucinolysome-encoding *Limousia* have low abundance in various human and animal gut microbiome samples. The seven human MAGs have a relative abundance between 0.06% and 0.61% in their original samples (**Figure 6A**), lower than those reported for *R. gnavus* (average 0.67% (*51*))*, A. muciniphila* (2.62% (*52, 53*)), and *C. perfringens* (average 1.16% (*52*)) in healthy humans. The relative abundances of *Limousia* in other animals are also generally lower than 1% (**Figure 6A** and **Figure 7D**). The prevalence of *Limousia* also exhibits large differences in different animals that have been studied: chicken (7.34%), goat (0.81%), and pig (19.63%). No other human microbiome samples beyond those listed in **Table 1** were mapped to *Limousia* MAGs, while the reported prevalence of *R. gnavus* in healthy humans is very high (average 43.1% (*51*)), as is *A. muciniphila* (47.1% (*52, 53*)) and *C. perfringens* (average 29.3% (*52*)). Nevertheless, it is of note that *Limousia* 16S RNAs were mapped to three human samples (one ancient human and two rural humans). A search using the ET540 16S RNA gene to query the NCBI NT database found another 16S RNA match (FJ503849) from an intestinal mucosal biopsy of a Crohn’s patient with a 99.6% sequence identity. All these suggest that, despite their very low abundance, the evidence is strong for the existence of mucinolysome-encoding *Limousia* in humans, as they are present in historically and geographically distanced samples.

To summarize, our findings reveal a previously unrecognized mechanism for mucin glycan degradation and provide a framework to investigate how microbial activity shapes gut health, influences the balance between mucin glycan degradation and mucin turnover, and affects host–microbe interactions and nutrient dynamics across diverse animal species.

## MATERIALS AND METHODS

### Search for mucinolysome-encoding MAGs in UHGG

We downloaded cohesin and dockerin hidden Markov models (HMMs) from Pfam V37.0. The length of the cohesin and dockerin profile HMMs were 139 and 58 residues respectively (*54, 55*). Using hmmsearch (http://hmmer.org/), we searched the dockerin and cohesin HMMs against the Unified Human Gastrointestinal Genome (UHGG) database (*21*) (**Figure 1A**). UHGG contains 289,232 genomes; most of them are MAGs and 3.8% are isolate genomes. In the 4,333 MAGs that have both dockerin and cohesin modules, we searched for mucin-degrading CAZyme domains (**Figure 2C**) in the Doc-containing proteins by using dbCAN3 (*56*) with default parameters.

### Expand the search for mucinolysome-encoding MAGs in other sources

To expand our search for mucinolysome-encoding MAGs, the scaffoldins from the human MAG MGYG000005496 (*57*) were used as queries to search against the RefSeq database and a total of 1,013,565 MAGs from various sources (**Table S8**, **Figure S6A**). The search was conducted using MMseqs2 (*58*) with sequence identity ≥ 50%, coverage ≥ 85%, and E-value < 1e-5. MAGs with at least one scaffoldin match were kept and further searched for the co-presence of cohesin and dockerin modules, and for mucin-degrading CAZymes as described above.

To identify peptidases in the MAGs, DIAMOND was used (E-value < 1e-10) (*59*) to search against the MEROPS database (*60*). Finally, sulfatases in the MAGs were identified using hmmsearch (http://hmmer.org/) (E-value < 1e-3) against the SulfAtlas database (*50*).

### MAG taxonomic classification, quality check and phylogenetic analysis

MAGs were classified taxonomically using GTDB-Tk (Genome Taxonomy Database, version 220) (*61*). CheckM2 v.1.0.2 (*62*) was used to assess the MAG completeness and contamination levels. To infer species phylogeny based on MAGs, we used Panaroo (*63*) to generate the single-copy core gene alignments, FastTree (*64*) for tree inference, and iTOL (*65*) for visualization. Inter-MAG average nucleotide identity (ANI) was calculated by FastANI (*66*) and ANI dendrogram was generated using Python scripts.

### MAG relative abundance and prevalence by read mapping of microbiome samples

For read mapping against MAGs in their original samples (**Table S9**), we used a pipeline (**Figure S6B**) consisting of bowtie2 (*67*), SAMtools (*68*), InStrain quick_profile (*49*) and CoverM (https://github.com/wwood/CoverM). Raw SRA files were quality-trimmed, and adapters and human contaminants were removed using bbduk from the BBTools suite (*69*), kraken (*70*) and Seqtk (https://github.com/lh3/seqtk). Remaining reads were normalized to 10 million using SeqKit (*71*), and quality was assessed using FastQC (*72*).

For read mapping against MAGs in 2,897 fecal samples of different animal hosts (**Table S10**), we used the same pipeline as above (**Figure S6B**). From the read mapping results, breadth of coverage (BC) and average read depth (ARD) were calculated for each MAG in each sample. BC of a MAG was calculated as the count of bases that have at least one read mapped divided by the count of all bases in the MAG (i.e., the length of the MAG). ARD of a MAG was calculated as the total lengths of all mapped reads divided by the length of the MAG. A MAG was considered present in a sample if it had a BC of ≥0.6 and ARD of ≥1. MAG relative abundance was calculated in each sample and MAG prevalence was calculated across all samples. MAG relative abundance was calculated as the count of the mapped reads to the MAG divided by the count of all reads in the sample. MAG prevalence was calculated as the number of positive samples divided by the total number of samples.

Read mapping was also performed against 16S RNAs and coding sequences (CDSs) of five scaffoldin proteins and 21 GHs of ET540. Proteinortho (*73*) was run on all 65 MAGs using parameters “-cov=90 -identity=80 -sim=1 -e=1e-30” to find protein orthologs of CDSs. The presence, BC and ARD of CDSs were determined in the same way as MAGs. 16S RNAs were predicted in 65 MAGs with Barrnap (https://github.com/tseemann/barrnap/tree/master). A 16S RNA was considered present in a sample if it had a BC of ≥0.6, ARD of ≥1 and read alignment sequence identity > 99% (inStrain_profile: --min_read_ani = 0.99). These thresholds are more stringent than those used in a recent study (*20*) to minimize false matches.

### Protein structure-based protein-protein interactions

We first extended all Pfam predicted dockerin and cohesin domain regions to 10 amino acids on both sides. We then extracted these extended sequences for structure prediction. For sequences of each Coh-Doc pair, we ran AlphaFold3 (AF3) (*43*) to model their complex structure and predict protein-protein interaction (PPI) interface. AF3 produced various scores to measure the quality of the predicted structure and PPI, such as pTM (predicted template modeling score), ipTM (interface-predicted TM-score), IF_contacts (number of interface contacts), and IF_pLDDT (average local distance difference test score) for the predicted interfacial residues. We further ran FoldDock (*44*) to calculate a pDockQ (predicted docking model quality) score for each predicted Coh-Doc complex structure from AF3. pDockQ score combines the IF_ pLDDT and IF_contacts scores. A pDockQ score >0.3 was used as the threshold to filter out Coh-Doc complex structures with low PPI prediction confidence.

### Growth media for *L. pullorum* ET540/kol109

The ET540 strain was routinely passaged on Columbia agar plates, which comprised Columbia broth (Neogen, Lansing, MI USA) solidified with 1.5% w/v agar (Franklin Lakes, NJ USA). Mucin-dependent growth was characterized in defined basal medium (DBM, **Table S6**) or defined basal medium, supplemented with 0.025% sodium sulfide and 0.2% tryptone (DBM #2) with no added carbon source, porcine gastric mucin (Type II mucin, Sigma Aldrich, St. Louis, MO USA), or with glucose added at the concentrations indicated in the figures. DBM agar plates were prepared by adding agar (BD) to 1.5% w/v. All growth experiments were performed at 37°C in an anaerobic chamber (Coy Laboratories, Grass Lake, MI USA) with an atmosphere of 5% H_2_, 5% CO_2_ and 90% N_2_. All media were pre-reduced in the anaerobic chamber for >24 hr.

### Initial validation of *L. pullorum* ET540

The ET540 strain was obtained from Dr. Ivan Rychlík (Veterinary Research Institute, Brno, Czech Republic). Glycerol stock of strain ET540 was streaked on Columbia agar plates and incubated at 37°C in an anaerobic chamber (gas atmosphere of 5% H_2_/5% CO_2_/90% N_2_) for 6-7 days to obtain colonies with distinct morphology. Colonies were re-passaged on Columbia agar in an anaerobic chamber to ensure purity, DNA was isolated for validation of strain identity by whole genome sequencing (details below), and new stocks in Columbia broth and preserved with 20% glycerol were created. For subsequent experiments, *L. pullorum* ET540 was passaged from frozen stock onto Columbia agar for 6-7 days prior to use.

### DNA sequencing and analysis of *L. pullorum* ET540

Full genome sequencing was performed by Plasmidsaurus using a combination of long-read sequencing with Oxford Nanopore and short-read Illumina sequencing. Long-read sequencing was performed on the PromethION P24 with R10.4.1 flow cell; libraries were prepared with the rapid barcoding kit 96 V14. The basecalling model was ont-doradod-for-promethion on super-accurate mode. Read quality was set for a minimum Qscore of 10 with adapters trimmed by MinKnow. Genomes were assembled by removing 5% of FASTQ reads with lowest quality scores with Filtlong v0.2.1 (https://github.com/rrwick/Filtlong), followed by downsampling the reads to 250 MB via Filtlong to create a rough assembly with Miniasm v0.3 (*74*). Reads were then re-downsampled to ∼100X coverage with heavy weight applied to remove low quality reads. High-quality ONT reads were assembled with Flye v2.9.1 (*75*) and polished with the reads generated during downsampling to ∼100X coverage using Medaka v1.8.0 (https://github.com/nanoporetech/medaka). Genes were annotated with Bakta v1.6.1 (*76*), contigs were analyzed with Bandage v0.8.1 (*77*), genome completeness and contamination were analyzed with CheckM v1.2.2 (*78*) and species/plasmids were identified through a combination of Mash v2.3 (*79*) against RefSeq genomes+plasmids, Sourmash v4.6.1 (*80*) against GenBank, and CheckM v1.2.2. Short-read Illumina sequencing was performed on a NextSeq12000 using a 2X150 bp read kit; libraries were prepared using SeqWell ExpressPlex 96 library prep kit. BWA (*81*) was used to align raw Illumina FASTQ reads to the ONT-only assembly (R1 and R2 aligned separately). BWA mem was used to align short reads to remove low-quality sequences from the ends of reads. Polypolish v0.6.0 (*82*) was used to generate the final polished FASTA sequence, which was then annotated with Bakta using default parameters.

### Characterization of mucin-dependent growth of *L. pullorum* ET540

To initially characterize mucin-dependent growth by ET540, colonies grown on Columbia agar plates were resuspended in Phosphate Buffered Saline (PBS), serially diluted in PBS to 10^-5^ dilution, and 100 μL of each dilution were spread on agar plates without added carbon source or with 0.2% w/v mucin under anaerobic conditions. Plates were incubated anaerobically at 37°C for 5–6 days, after which the number of colony forming units (CFU) on the plate with >50 colonies and <250 colonies was enumerated to determine CFU/mL; based on dilutions, the limit of detection was <10 CFU/mL. Images of plates were captured using Syngene NuGenius, and all images were adjusted to the same brightness and contrast in Adobe Photoshop prior to embedding in Adobe Illustrator.

Maximum cell densities (OD_620_) and growth rates were determined for cultures grown in DBM or DBM #2 broth with added glucose or mucin. Colonies grown on Columbia agar were used to inoculate DBM + 0.2% mucin broth cultures, which were allowed to grow anaerobically at 37°C for 24-48 hrs. Cultures were then diluted back to 5% v/v into broth with no carbon source, or with different concentrations of mucin or glucose in 96-well culture plates. Growth was monitored over time by measurement of OD_620_ in a Tecan Sunrise plate reader (Tecan, Männedorf, Switzerland) with measurements collected every 20 minutes for 36-72 hr. OD_620_ data for each replicate well (n ≥ 3/condition) were normalized to the starting OD_620_. Growth rate (μ) was determined by linear regression of natural log transformed OD_620_ data during exponential growth (2-24 hr). Doubling times were calculated from growth rates (ln(2)/μ). Data were plotted in GraphPad Prism v10.5.0.

### Growth of ET540 bacterial strain for RNA sequencing

To evaluate transcriptomic responses of ET540 in response to mucin or glucose as carbon sources, triplicate cultures were grown in 40 mL DBM #2 with 0.2% mucin for 48 hr. Cultures were then diluted to 5% v/v into either 100 mL DBM #2 with 0.2% mucin or 0.2% glucose in triplicate for collection of samples for RNA extraction. In parallel, inoculum was diluted back to 5% v/v into 96-well plates containing the same growth media, and growth was monitored in a Tecan Sunrise plate reader as described above. Samples (50 mL) were collected for RNA extraction after 12 hr of growth (late exponential phase based upon growth of parallel cultures grown in plate reader), pelleted by centrifugation at 4,800 × g for 15 min at 15°C, decanted and resuspended in 1 mL of TRIzol reagent (Invitrogen, Waltham, MA USA); samples were stored at −80°C until extraction.

### RNA extraction

RNA was extracted and purified using the Trizol extraction protocol with Qiagen RNeasy. Specifically, 0.2 mL of chloroform per 1 mL TRIzol reagent was added. The tubes were vigorously shaken for 15 sec, incubated at room temperature for 2–3 min, and centrifuged at 12,000 × g for 15 min at 4°C. The upper aqueous phase (approximately 60% of the original TRIzol reagent volume) was carefully transferred to a fresh RNase-free tube, and an equal volume of 75% ethanol was added and mixed gently. The RNA-ethanol mixture was transferred onto a RNeasy spin column and centrifuged at ≥8000 × g for 15 sec at 4°C. Subsequent steps were performed according to the manufacturer’s protocol, with elution in a total of 80 μL of RNAse-free water (two-step elution with 50 μL and 30 μL).

The extracted RNA was quantified using a NanoDrop spectrophotometer (Thermo Fisher Scientific), assessing concentration, total RNA yield, and purity (A260/A280 and A260/A230 ratios). RNA integrity was assessed with a 4200 Agilent TapeStation following manufacturer’s protocols, ensuring that high-quality standards required for RNA sequencing were met. RNA samples were aliquoted into RNase-free tubes and stored at −80°C until shipment to the sequencing facility.

### RNA-seq differential gene expression analysis

Paired-read FASTQ files from two groups, control group G and mucin group M (each with 3 replicates) were analyzed for differential gene expression. The read files were quality assessed using FastQC (*72*), aggregated using MultiQC (*83*) and quality cleaned with trimmomatic (*84*). Post trimmed files were quality assessed again with FastQC. Using the reference-based method, HISAT2 (*85*) and samtools (*68*) were used to align clean reads against the ET540 reference genome and generate BAM files. We used gffreads (*86*) to convert ET540 genome to “.gtf” format. The featureCounts program (*87*) was then used for the quantification step to count the reads. Using the DESeq2 (*48*), differentially expressed genes between the control group (G) and the mucin group (M) were identified using an adjusted p-value (FDR) threshold of < 0.05 and an absolute log2-fold change (∣log2FC∣) threshold of > 1 (2-fold change). KEGG (*88*) pathway enrichment analysis was performed using clusterProfiler v4.12.6 (*89*).

## ACKNOWLEDGEMENTS

This work was partially completed utilizing the Holland Computing Center of the University of Nebraska—Lincoln.

## FUNDING

This work was supported by the National Institutes of Health (NIH) awards [R01GM140370 and R03OD039979], and partially by the United States Department of Agriculture (USDA) award [58-8042-3-076], a Nebraska Research Collaboration Initiative Award, and the Nebraska Tobacco Settlement Biomedical Research Enhancement Funds. Funding for open access charge: R03OD039979. Thanh Do was supported by an assistantship from the Department of Food Science and Technology and the Agricultural Research Division of UNL.

### Conflict of interest statement

None declared.

### Data availability statement

The raw sequencing reads of ET540 genome are available at https://www.ncbi.nlm.nih.gov/bioproject/?term=PRJNA1339042. The faa, fna and gff files of all 65 mucinolysome-encoding *Limousia* genomes have been deposited in figshare at https://doi.org/10.6084/m9.figshare.30192742.

### Code availability statement

Code is freely available at https://github.com/bcb-unl.

## Supplementary figures

**Figure S1.**
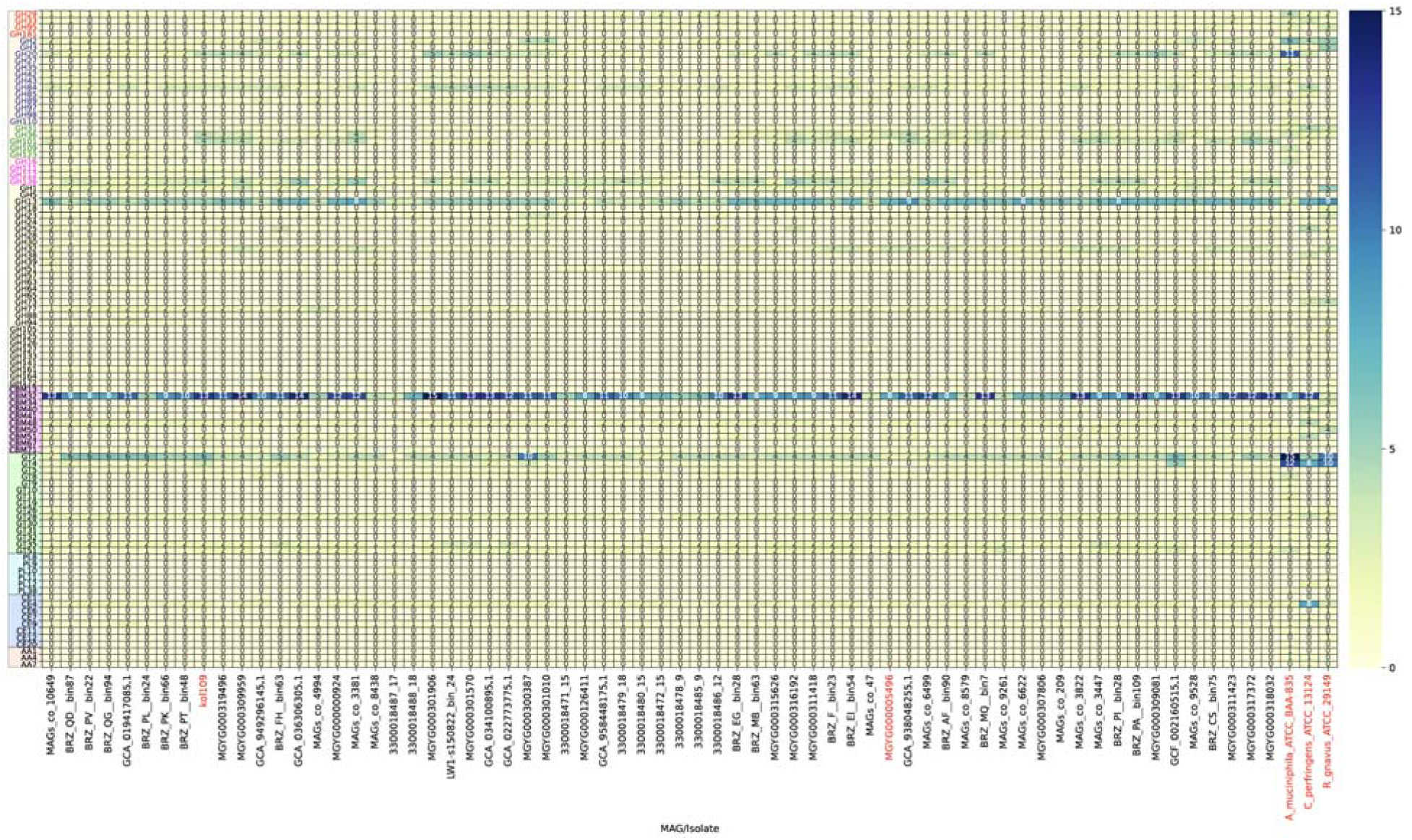
Heatmap of all CAZyme families (rows) in 65 Limousia genomes and three known mucin-degrading genomes (columns). The values indicate the number of genes per bacterium for the given protein family. Colored GH families are known for mucin degradation according to the literature.

**Figure S2.**
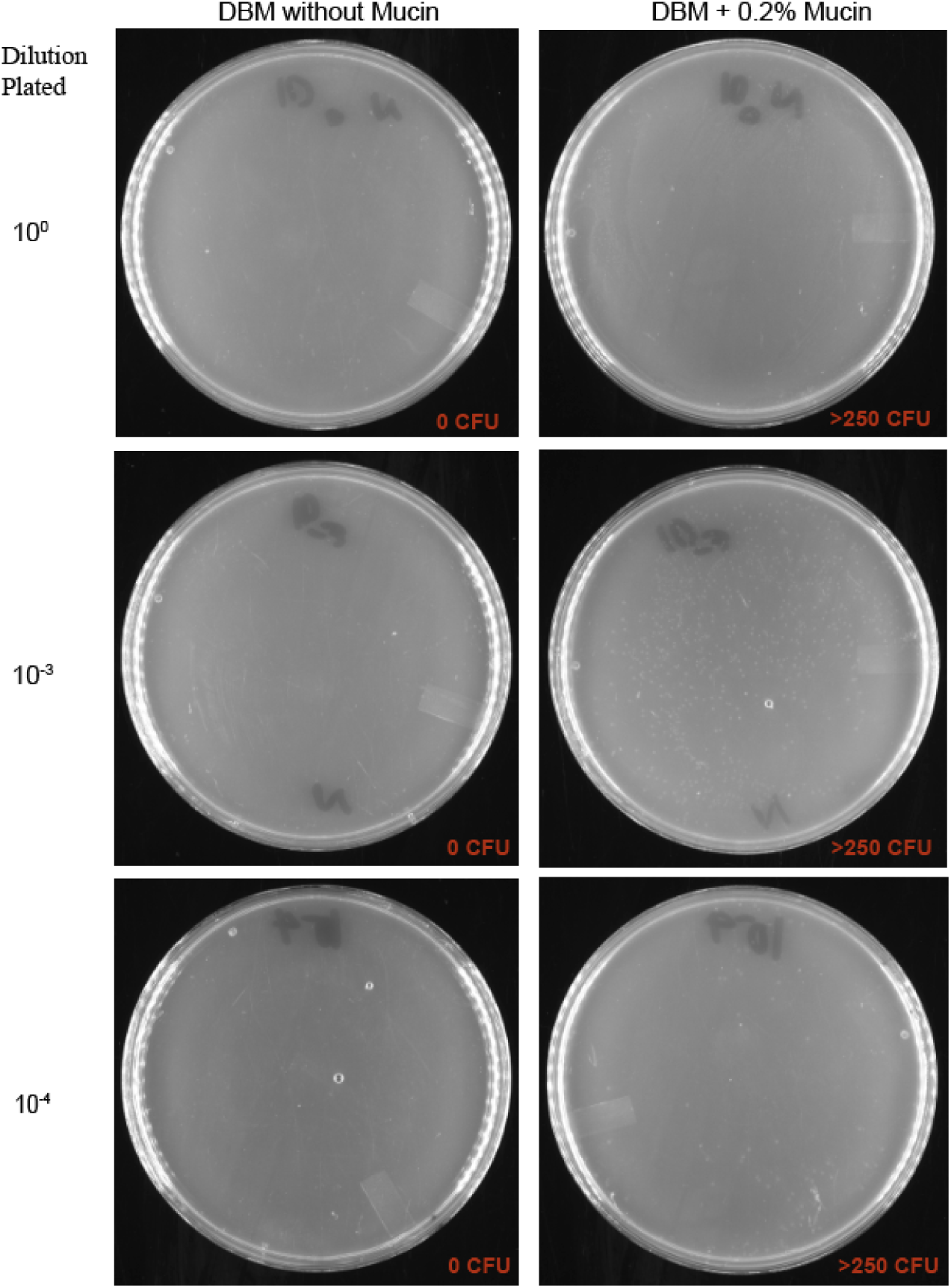
Limousia pullorum ET540 grows on DBM with 0.2% mucin as the sole carbon source. Growth of L. pullorum ET540 on DBM, without an added carbon source or with 0.2% porcine gastric mucin added as the sole carbon source, was assessed by measuring colonies formed from L. pullorum at different dilutions of inoculum. Colony forming units (CFU)/mL were calculated as described in methods, with DBM without mucin allowing recovery of <10 CFU/mL and DBM + 0.2% mucin allowing recovery of 6.1 × 10^6^ CFU/mL.

**Figure S3.**
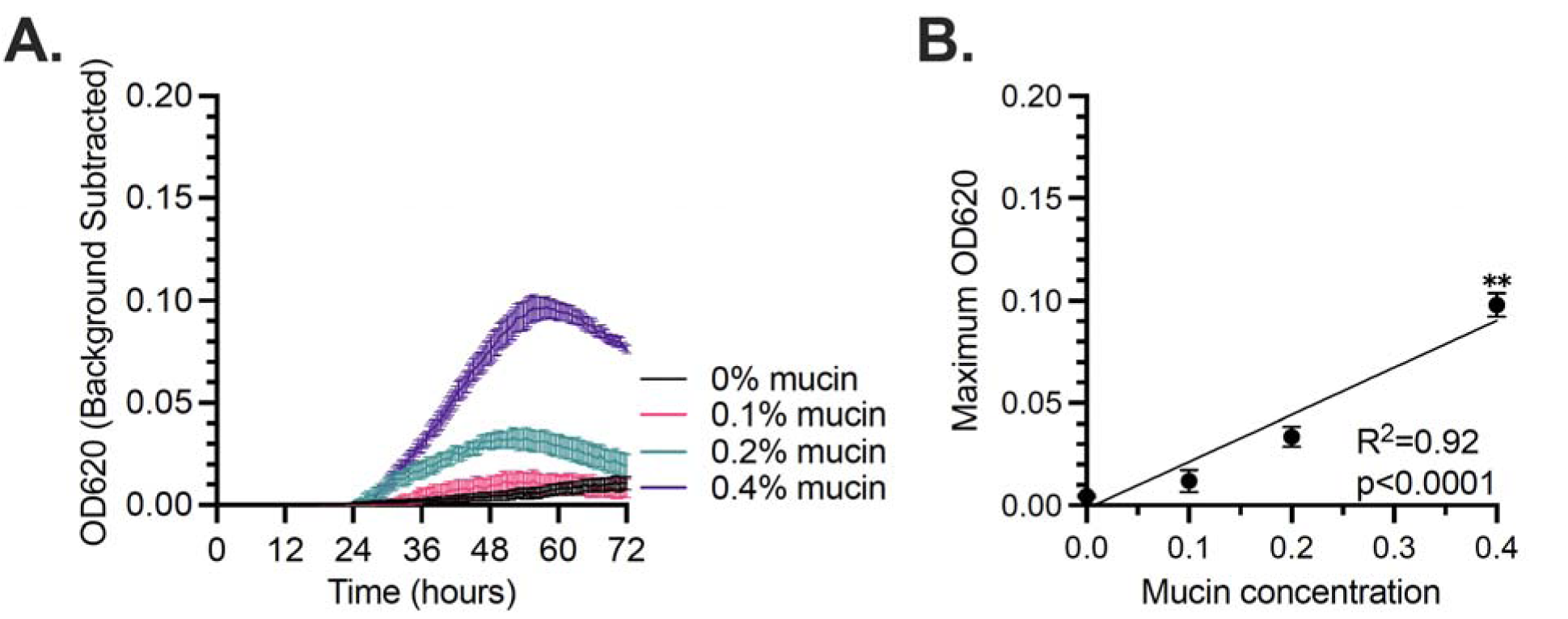
Repeat growth of L. pullorum ET540 in DBM + mucin. L. pullorum ET540 was grown in DBM + 0.2% mucin for 24 hr, diluted 5% v/v in fresh DBM containing the indicated mucin concentrations, and (**A**) growth over time in culture and (B) maximum OD_620_ relative to mucin concentration were plotted. In (A), mean ± SEM were plotted for triplicate cultures at each time point. In (**B**), mean ± SEM were plotted for triplicate cultures at each mucin concentration. Linear regression was used to determine the goodness of fit (R2) and significance of slope deviation from zero, with p<0.05 reported. Significance of differences in maximum OD_620_ values at each mucin concentration relative to no mucin was determined by one-way ANOVA with Brown-Forsythe correction for unequal variances and Dunnett’s T3 correction for multiple comparisons. **, p<0.01.

**Figure S4.**
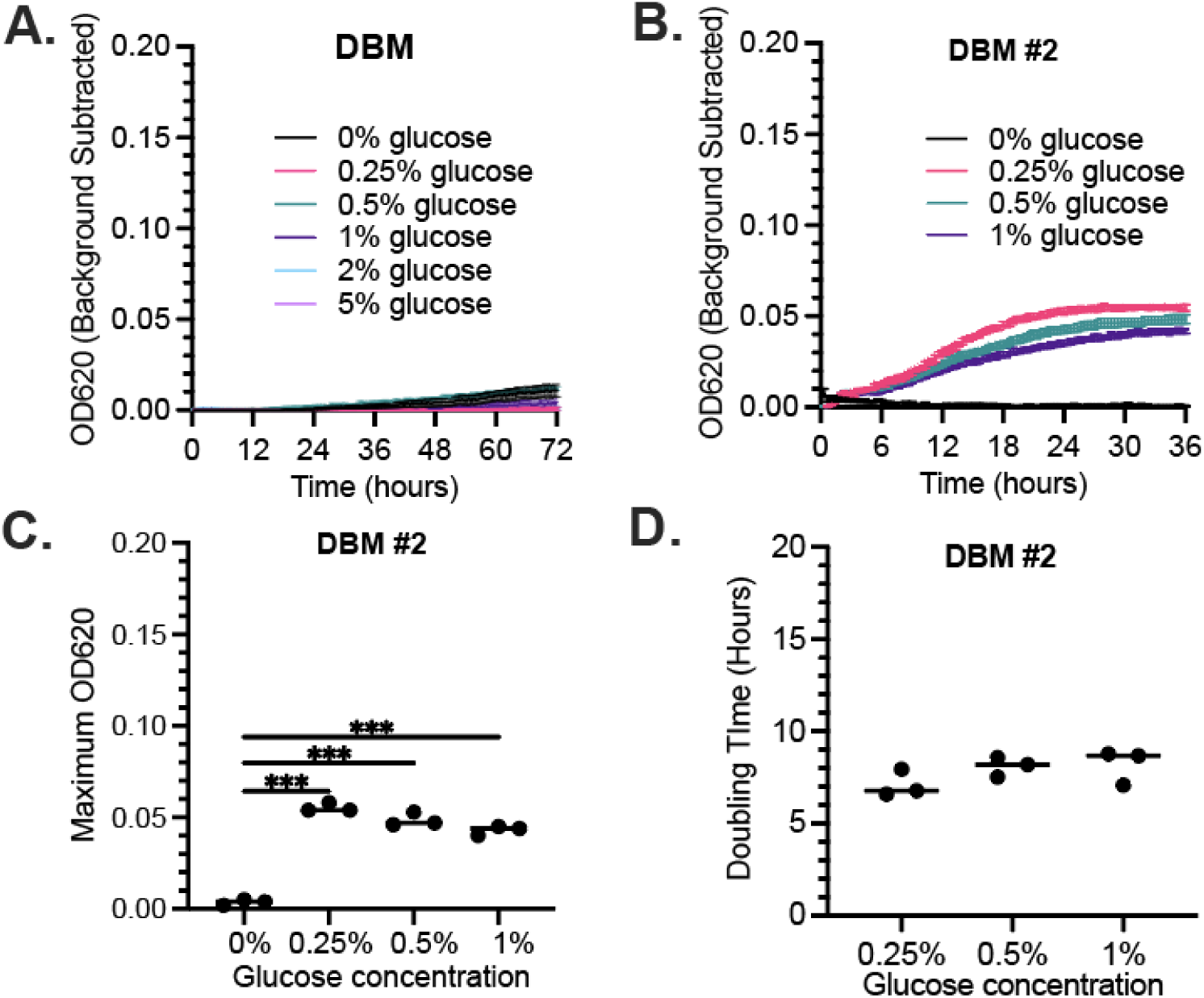
Growth of L. pullorum ET540 in defined medium with glucose as the sole carbon source. (**A**) L. pullorum ET540 was grown in DBM + 0.2% mucin for 24 hr, diluted 5% v/v in fresh DBM containing the indicated glucose concentrations, and mean ± SEM was plotted for triplicate cultures at each time point. In (**B**)-(**D**), L. pullorum ET540 was grown in DBM + 0.2% mucin for 48 hr, diluted 5% v/v in fresh DBM #2 containing the indicated glucose concentrations and (**B**) growth over time in culture, (**C**) maximum OD_620_ relative to glucose concentration, and (**C**) doubling time from 2-24 hr for each glucose concentration were plotted. In (**A**) and (**B**), mean ± SEM were plotted for triplicate cultures at each time point. In (**C**) and (**D**), each point represents a replicate with the line indicating the median. Significance of difference in maximum OD_620_ from no added glucose and doubling time between mucin concentrations was determined by one-way ANOVA with Brown-Forsythe correction for unequal variances and Dunnett’s T3 correction for multiple comparisons. ***, p<0.001.

**Figure S5:**
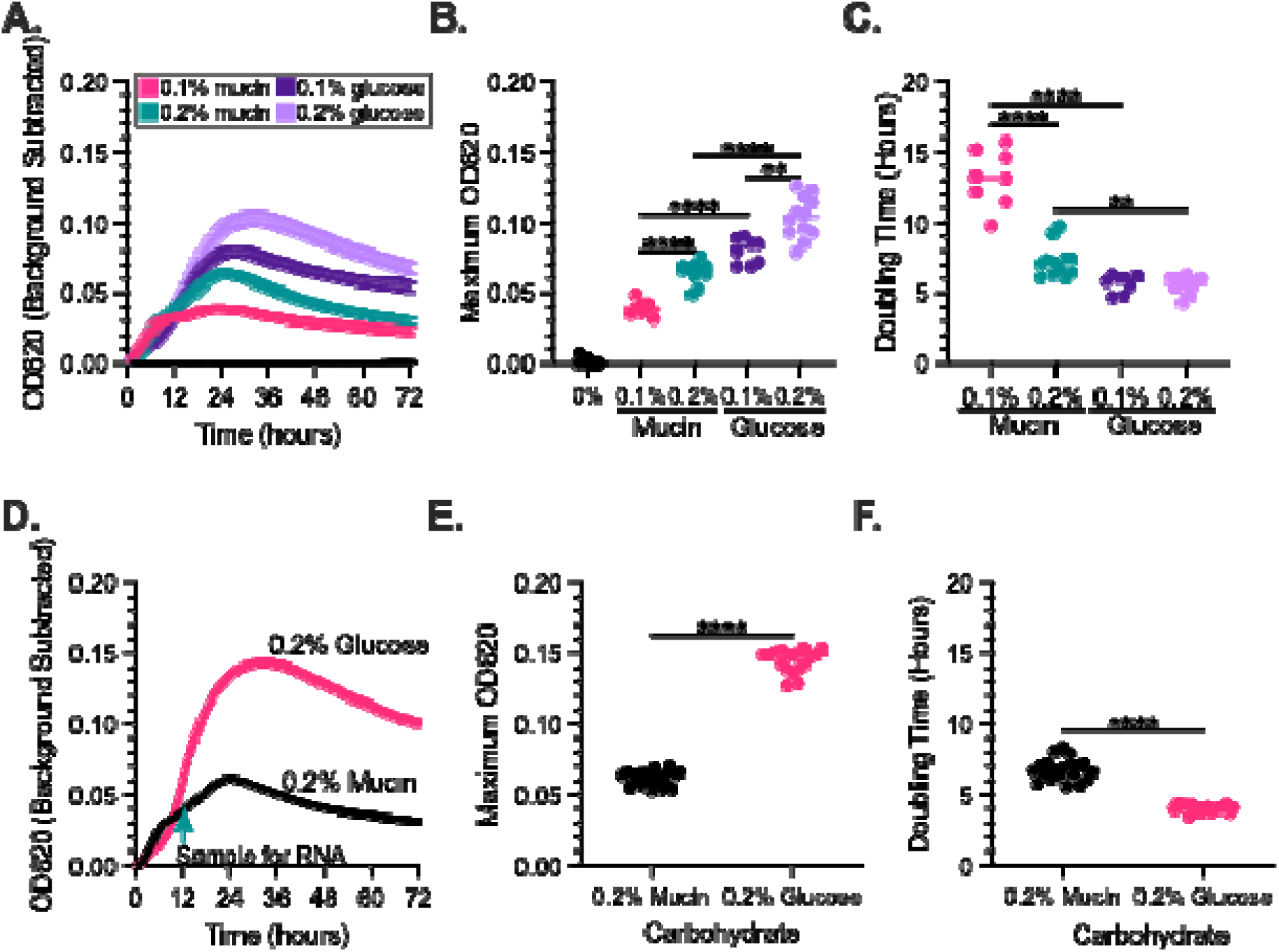
Growth of L. pullorum strains in modified defined basal medium yields growth with mucin or glucose. In (**A**)-(**C**), L. pullorum ET540 was grown in DBM + 0.2% mucin for 48 hr, diluted into fresh DBM #2 medium with 0.1% or 0.2% mucin or glucose, and (**A**) growth over time in culture, (**B**) maximum OD_620_ relative to mucin concentration, and (**C**) doubling time from 2-24 hr for each mucin or glucose concentration were plotted. In (**A**), mean ± SEM values were plotted for replicate cultures at each time point. In (**B**), significance of differences in maximum OD_620_ between 0% and 0.1% or 0.2% of mucin or glucose, between 0.1% and 0.2% of the same substrate, or between the same percentage of different substrates (e.g., 0.1% mucin and 0.1% glucose), was determined by one-way ANOVA with Brown-Forsythe correction for unequal variances and Dunnett’s T3 correction for multiple comparisons. All cultures grown in medium with mucin or glucose were significantly higher than with no added carbohydrate (p<0.0001, not shown). In (**C**), significance of differences in doubling times between 0.1% and 0.2% of the same substrate or between the same percentage of different substrates was determined by one-way ANOVA with Brown-Forsythe correction for unequal variances and Dunnett’s T3 correction for multiple comparisons. In (**D**)-(**F**), L. pullorum was grown under the same conditions as in (**A**)-(**C**) in 0.2% mucin or 0.2% glucose to collect samples for transcriptomics. In (**D**), growth over time in culture was plotted as the mean ± SEM and the time of sample collection for RNA extraction is indicated by an arrow. In (**E**) and (**F**), each point represents a replicate with the line indicating the median. Significance of differences in maximum OD_620_ or doubling time between cells grown in mucin or glucose was determined by two-tailed student’s t-test with Welch’s correction for unequal variances.

**Figure S6:**
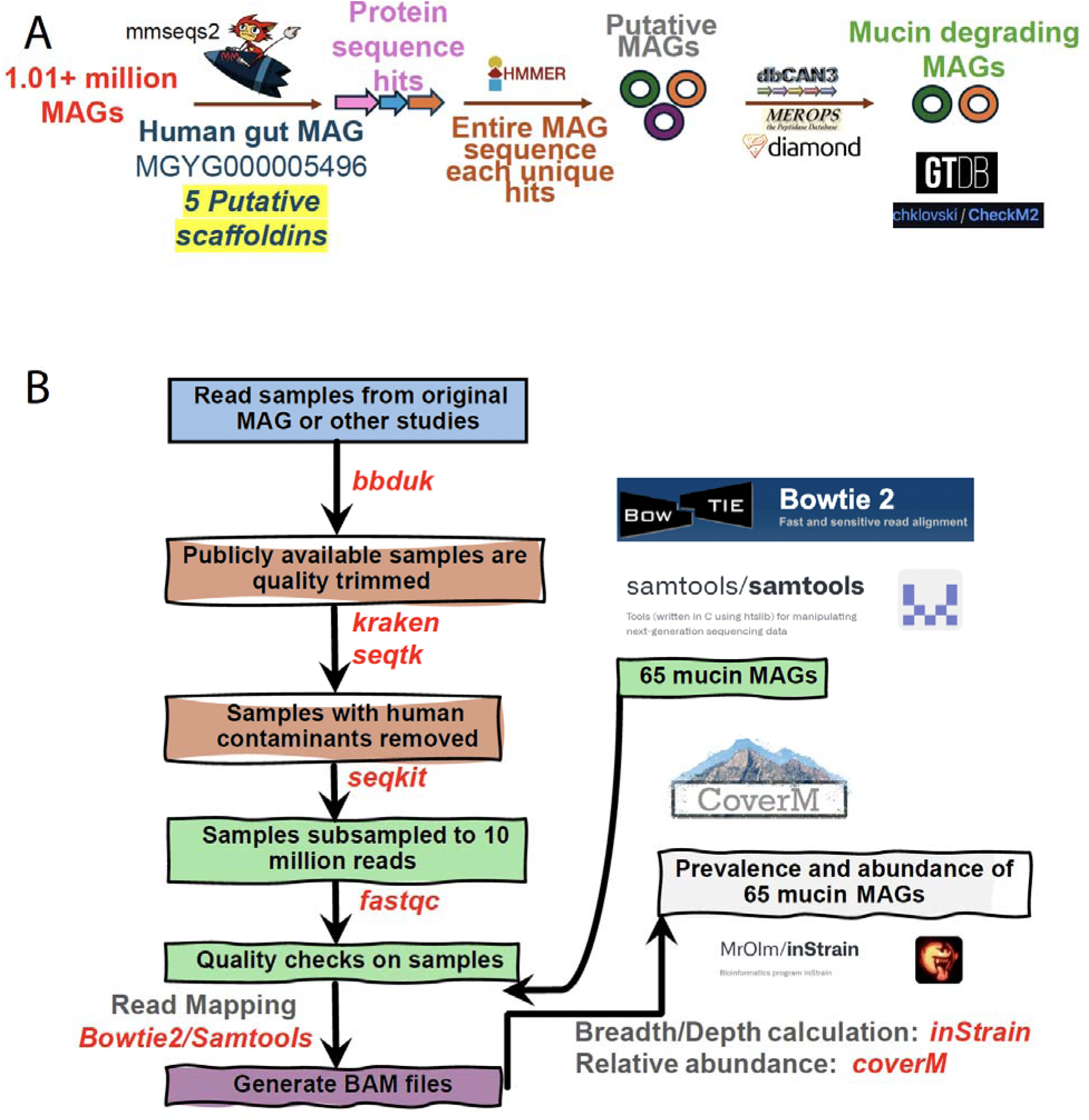
Computational workflows and tools. (**A**) Identification of the 65 MAGs. (**B**) and Read mapping to determine MAG abundances in their original samples and MAG prevalence in 2,897 fecal samples of different human and animal hosts.

## Supplementary tables

**Table S1: 65 genomes that encode mucinolysomes**

**Table S2: Functional domains in proteins of mucinolysomes of 65 genomes**

**Table S3. CAZymes in five genomes**

**Table S4: PPIs predicted for Coh-Doc pairs in *C.* perfringens ATCC 13124**

**Table S5: PPIs predicted for Coh-Doc pairs in Lpuc ET540/kol109**

**Table S6: Chemically defined media composition**

**Table S7: Gene expression of all genes in ET540**

**Table S8: All MAGs searched in this study**

**Table S9: SRA samples of 65 genomes**

**Table S10: 2,897 fecal samples of different animal hosts**

## Notes

https://doi.org/10.6084/m9.figshare.30340819.v1

### Competing Interest Statement

The authors have declared no competing interest.

### Summary of Updates

A co-author was in the PDF, but not in the HTML. I must have forgot to include this person when I submit it. Now I added this person in the submission system.

