## Supplementary figures and images for "Mucinolysome in gut microbiomes of farm animals and humans"

### Figure S1

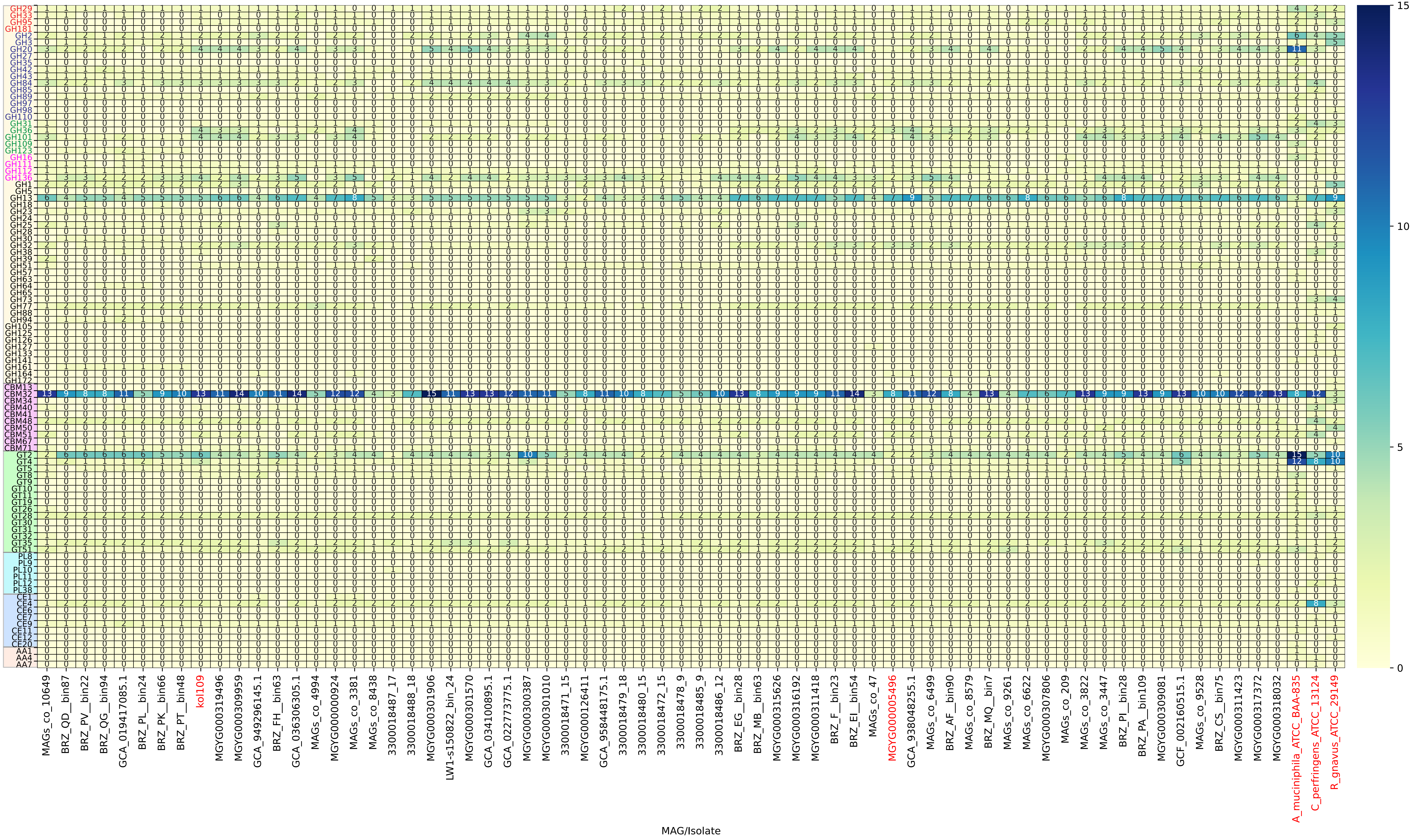

### Figure S2

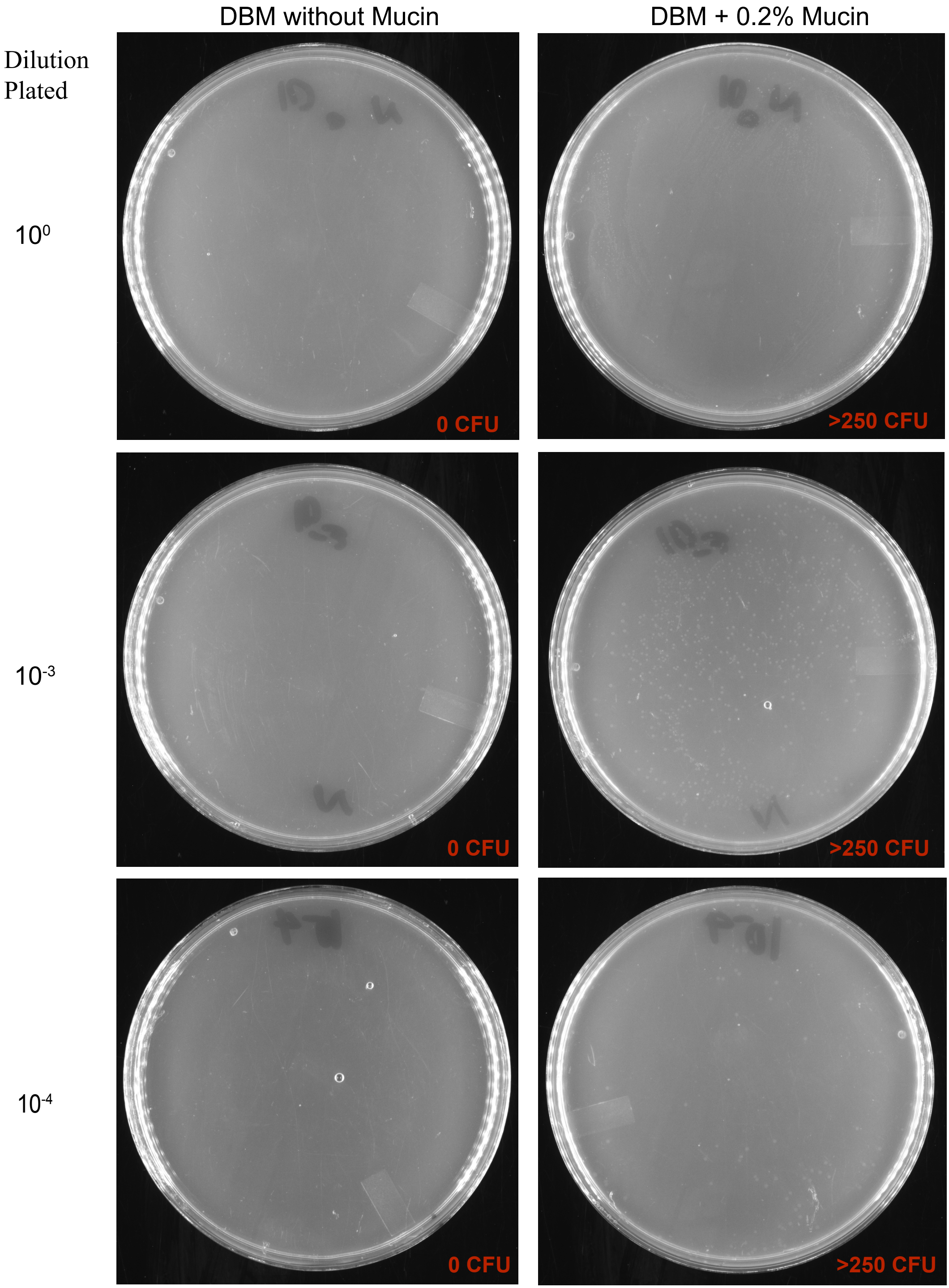

### Figure S3

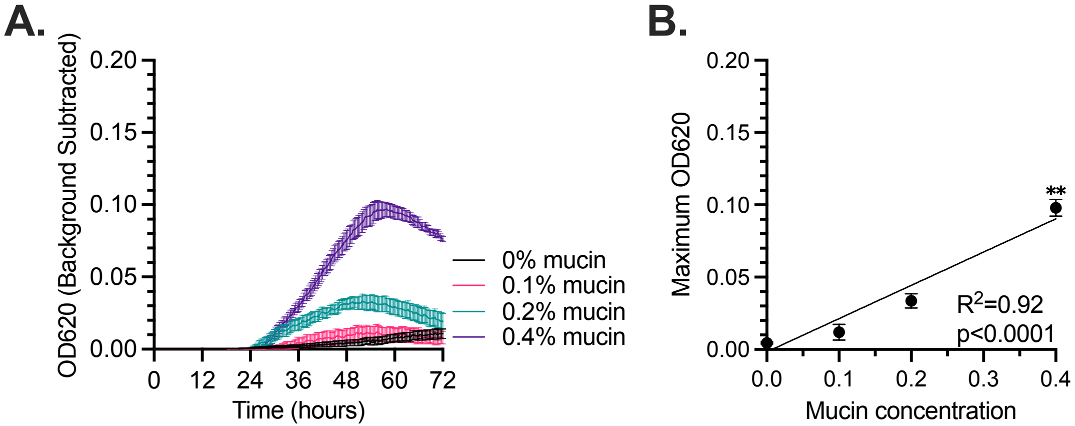

### Figure S4

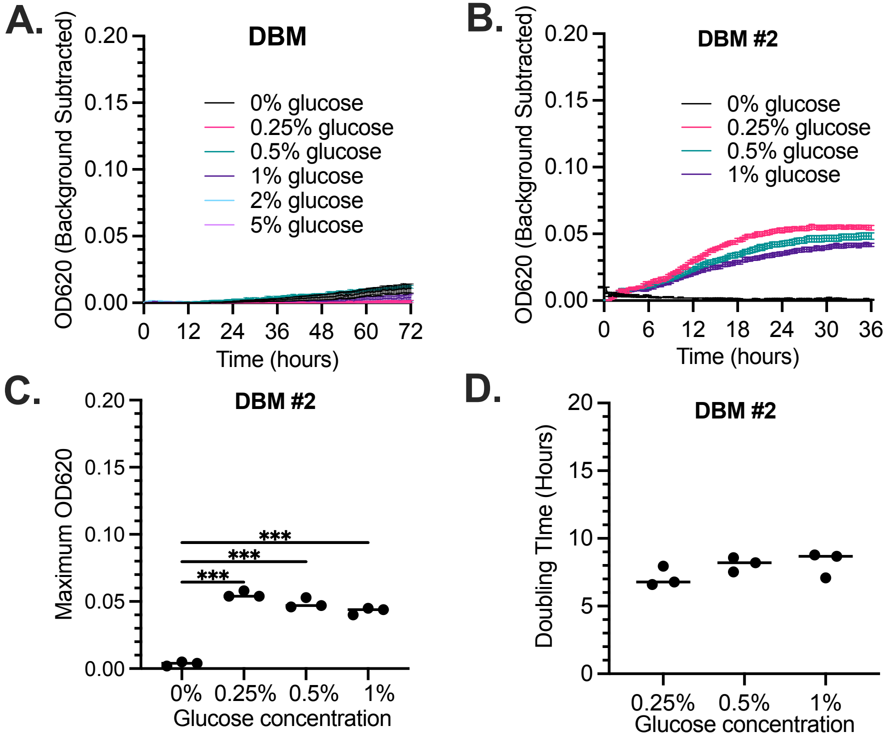

### Figure S5

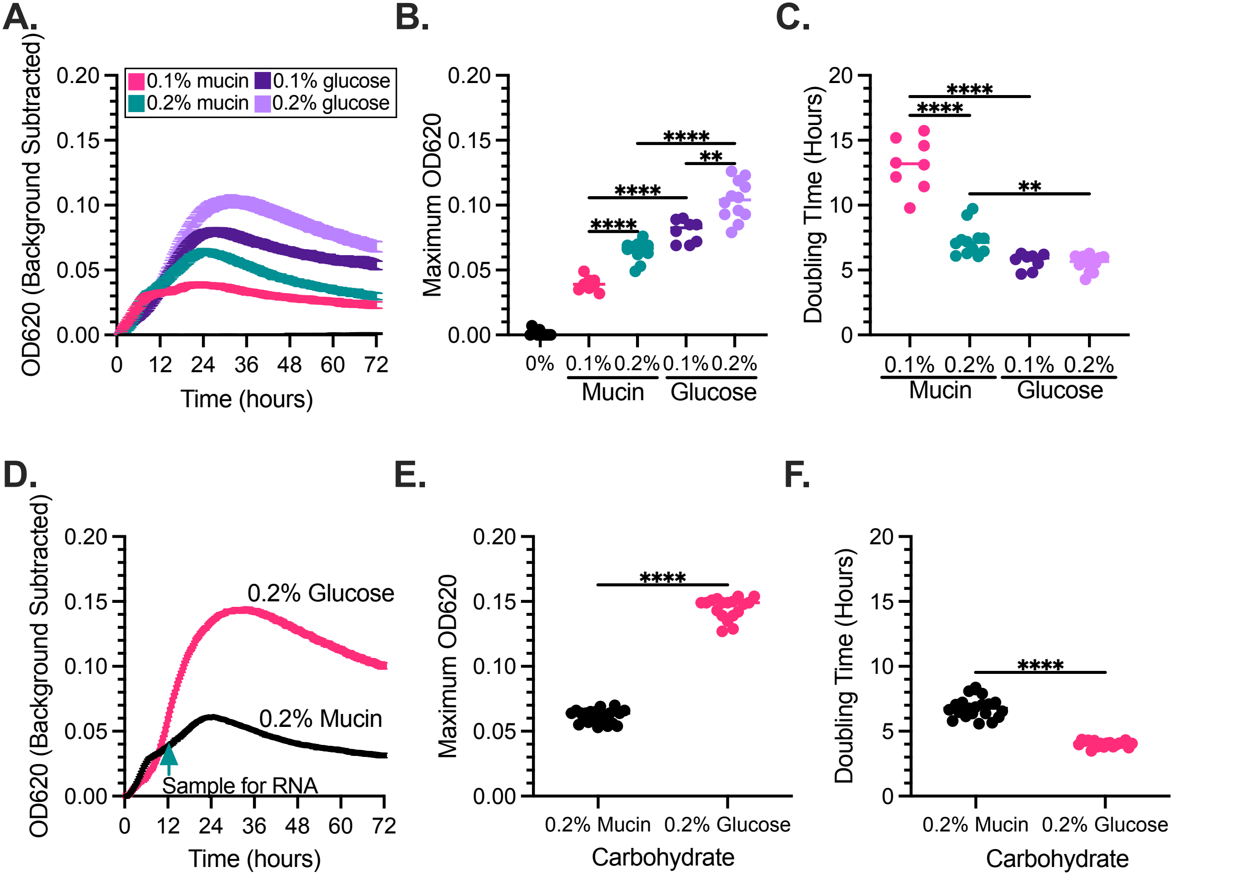

### Figure S6

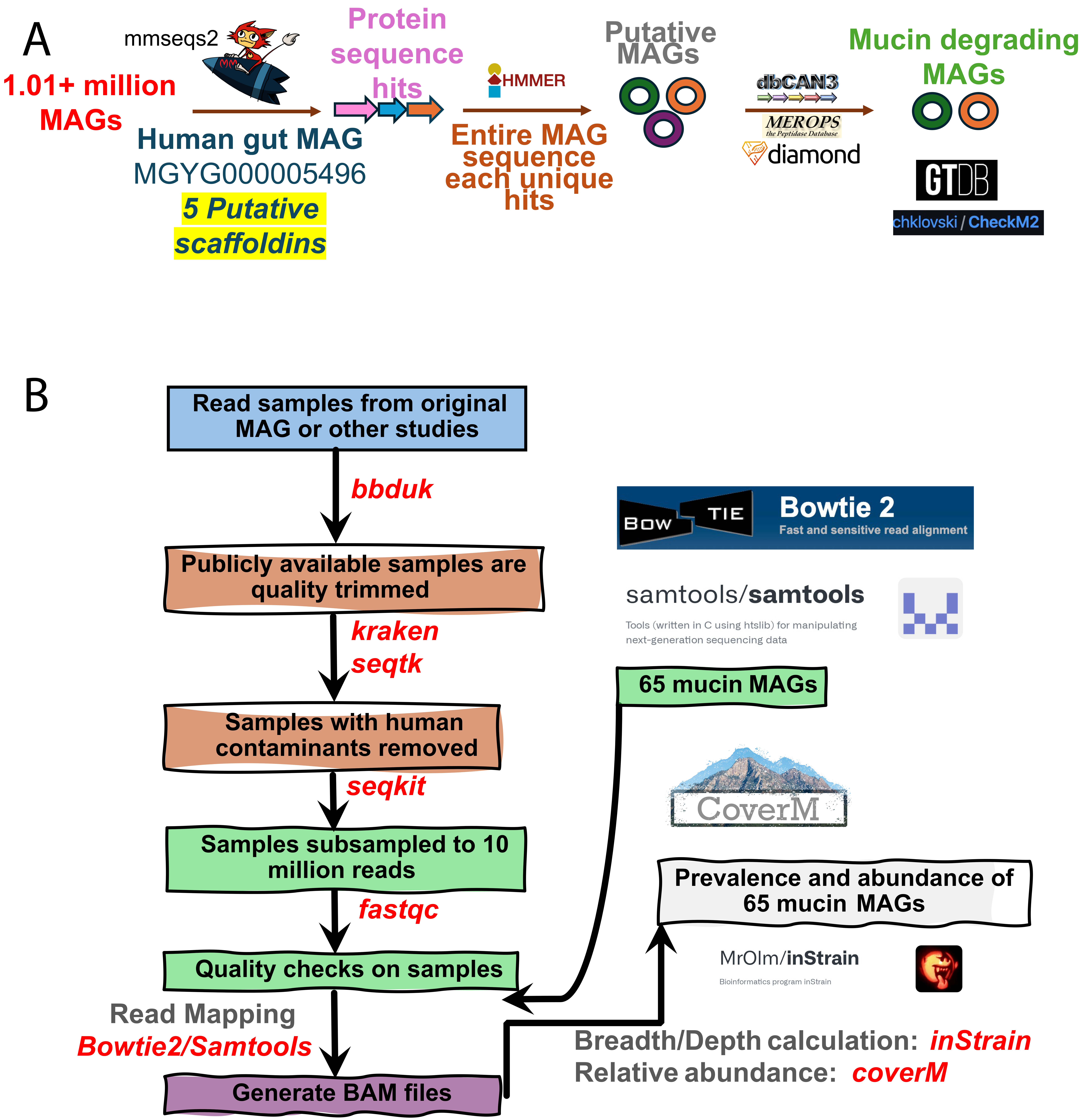
